# Uncovering the Bromodomain Interactome using Site-Specific Azide-Acetyllysine Photochemistry, Proteomic Profiling and Structural Characterization

**DOI:** 10.1101/2021.07.28.453719

**Authors:** Shana Wagner, Elena Fedorov, Babu Sudhamalla, Hum Nath Jnawali, Ronald Debiec, Agnidipta Ghosh, Kabirul Islam

## Abstract

Protein-protein interactions mediated by acetyllysines and bromodomains are essential for the regulation of eukaryotic gene expression. Diagramming the bromodomain interactome network and molecular characterization of these interactions are key to unraveling context-dependent signaling pathways. Herein, we employ a chemoproteomic platform, called interaction-based protein profiling (IBPP), to map the acetylome of the bromodomain and extra terminal domain (BET) family of bromodomains. We developed photo-responsive bromodomain analogues to carry out UV light-induced azide-acetyllysine crosslinking within the recognition cavity of the domain to capture transient interacting partners present in human cells. Subsequent proteomic and biochemical analyses lead to the identification of an array of acetylated interacting factors, which extend the potential function of BET family proteins beyond transcription. We present here eight high-resolution crystal structures of interactome-bound bromodomains that underscore the atypical binding modes and sequence motifs by which BET members recognize an expanded repertoire of acetylated proteins. In addition, we report an acetyllysine-dependent interaction between BRD4 and interleukin enhancer binding factor 3 (ILF3) in human cells, and uncover its role in recruiting transcriptional regulators to the promoter of the Survivin gene, an anti-apoptotic factor overexpressed in a wide range of human neoplasia. Collectively, our work provides a blueprint for engineering bromodomains, establishes IBPP as a robust chemoproteomic tool to characterize bromodomain interactome, and reveals distinctive recognition modes by which such associations may take place in order to modulate signaling pathways. Furthermore, these structural snapshots offer clues to design specific small-molecule inhibitors against the novel BET interactions and paves the path for exploring the biological significance of this newly identified signaling network.

## Introduction

Bromodomains are interacting modules consist of approximately 110 amino acids, which recognize acetylated lysine in histones through its ‘hydrophobic cage’ to recruit transcriptional complexes for modulating gene expression (Fig. 1A, B).^1^ The human genome encodes 61 bromodomains in its 46 proteins which include transcriptional coactivators, chromatin remodelers, helicases and members of the bromodomain and extra terminal domain (BET) family.^2^ It is, however, becoming increasingly apparent that bromodomains interact with a wide range of acetylated non-histone proteins to diversify its biological footprint beyond transcription.^3,4^ How a large ensemble of these acetyllysine ‘readers’ function in chromatin and non-chromatin contexts is largely unexplored, primarily because a lack of systematic dissection of their distinct interactome. Exhaustive profiling of these interacting partners by traditional enrichment methods is challenging, as bromodomains display weak and dynamic binding towards acetylated proteins with dissociation constants (*K*_d_) typically ranging between 10 micromolar to 1 millimolar.^5^

**Figure 1.**
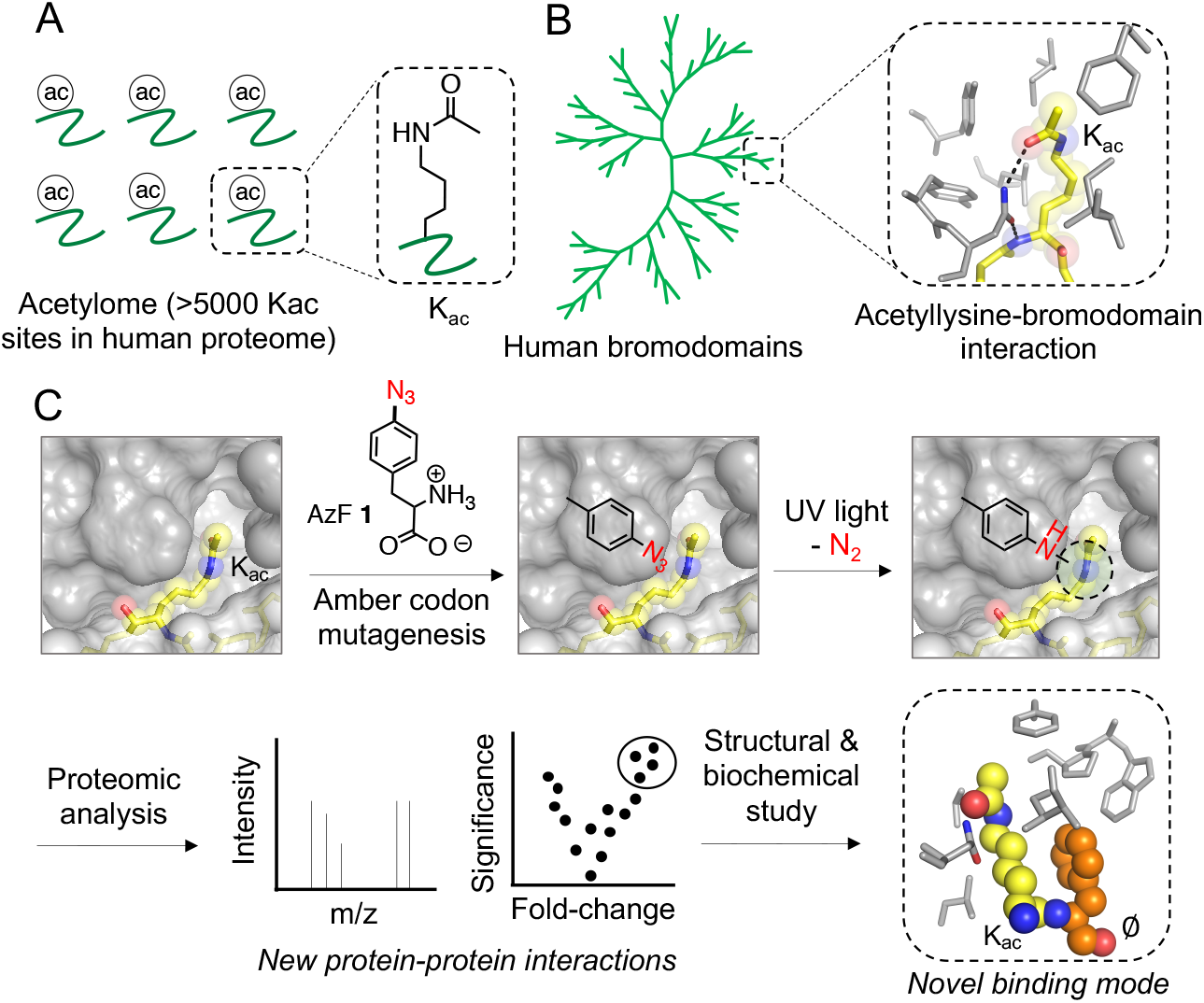
Schematic representation of IBPP for acetylome profiling. (A) Chemical structure of acetyllysine (Kac). The human proteome is estimated to have >5000 Kac sites. (B) Bromodomain family tree comprised of 61 bromodomains in 46 human proteins. Closeup of conserved Kac interaction (PDB 3uvy) in the hydrophobic pocket, i. e., ‘aromatic cage’ in a bromodomain. (C) Steps detail IBPP approach: (i) Amber suppressor mutagenesis to incorporate *p*-azido phenylalanine (AzF) **1** into bromodomain aromatic cage without altering its affinity towards Kac in proteins. (ii) photo-crosslinking to potential interacting partners for affinity enrichment, (iii) proteomic analysis, and (iv) biochemical and structural validation.

Members of the BET family, BRD2, 3, 4 and BRDT, each feature tandem bromodomains, and have emerged as critical regulators of gene transcription, cell cycle, and organogenesis; their aberrant expression is associated with inflammation, obesity, and cancer.^6^ BRD2 and 4 are essential for embryonic development, and phenotypes associated with their haploinsufficiency or null mutation cannot be rescued by other BET members despite high sequence homology and structural conservation.^7^ We reason that such non-redundant functions likely stem from differences in non-histone binding partners of individual BET proteins, which are known to recognize acetylated GATA1, RelA, Twist and MTHFD1 for controlling transcriptional and metabolic processes.^8–11^ Thus, it is imperative to delineate the acetyllysine interactome network for deciphering distinctive or partially overlapping functions within the BET proteins.

Chemoproteomic approaches have made a significant contribution towards cataloguing novel enzymes and their cognate substrates.^12^ However, it has remained a major challenge to capture the transient protein-protein interactions that drive the majority of the signaling cascades. This is often due to the absence of a catalytically active nucleophilic amino acid at the interface that is necessary to harness activity- and/or affinity-based chemoproteomic probes. Strategies involving proximity-induced covalent capture of interacting interfaces using reactive molecules such as disuccinimidyl sulfoxide (DSSO) are being developed.^13,14^ Such approaches, however, require precise positioning of nucleophilic amino acids in the interacting partners and often lead to an unproductive intra-link within the protein. Immunoprecipitation-based methods are only appropriate for identifying strong and abundant interactions and lack the necessary temporal control for profiling a dynamic interactome. Furthermore, such methods pose limitations in recognizing domain-specific binding partners. This is important, as a majority of the human proteins relay signals by employing specific domains such as the bromodomain. Recently, chemically customized nucleosomes and biotinylated readers have been developed to identify interacting proteins for a given histone modification.^15,16^ Although these tools enable characterization of the chromatin-dependent local interactome, we lack a method for unbiased profiling of proteome-wide non-histone interacting partners of a specific reader domain. Given that wild type readers employed in the earlier studies are often limited to stronger, multi-valent interactions, we reasoned that engineered readers with covalent binding capability would allow an entire interactome to be analyzed in a conditional and temporal manner.

Proteins and oligonucleotides carrying photoactive building blocks have been developed to capture dynamic interactomes.^17^ A particularly important approach includes site-specific introduction of a light- or chemical-induced crosslinker by read-through of the amber stop codon using an evolved aminoacyl synthetase-tRNA pair.^18,19^ Employing this strategy, we recently developed an interaction-based protein profiling (IBPP) platform to characterize the binding partners of BRD4 (Fig. 1C).^20^ By placing a photo-sensitive amino acid in the ‘aromatic cage’ of BRD4 using amber suppression mutagenesis, the engineered reader permitted covalent bond formation with dynamic interactome upon UV irradiation and facilitated enrichment of BRD4 complexes and subsequent proteomic analyses. The IBPP platform offers certain advantages over earlier approaches: i) site-specific installment of a photo-crosslinkable amino acid (PCAA) to profile only the acetylated interactome with temporal control using light, ii) ability to capture weaker interactions by forming a stable covalent bond, and iii) minimization of non-specificity due to precise crosslinking within the aromatic cage. Herein, we characterize the acetyllysine interactome of the BET members, BRD2, 3, and BRDT in particular, by employing the IBPP platform. We access photo-sensitive analogues of the readers and uncover their distinct as well as shared binding partners from human cells by proteomic analysis of the crosslinked species and subsequent validation of the putative targets. In addition, we report eight high-resolution crystal structures of bromodomains bound to novel acetylated non-histone protein segments identified by the IBPP platform detailed in this work. These structures reveal both canonical and unprecedented modes of binding, rationalizing how BET members recognize an extended repertoire of interacting partners in order to regulate a diverse set of cellular processes. In subsequent biochemical experiments, we validated the interaction between acetylated interleukin enhancer binding factor 3 (ILF3) and BRD4 in human cells. We demonstrated that acetyllysine-mediated ILF3-BRD4 interaction is critical for the transcriptional regulators to localize to the gene promoters including Survivin, a key mitotic and anti-apoptotic protein that is overexpressed in transformed cells.

## Results and discussion

### Designing BET bromodomain analogues

In spite of sequence divergence, bromodomains contain a highly conserved hydrophobic pocket, called an ‘aromatic cage’, to accommodate the acetylated lysines present in interacting partners (Fig. 1B).^1^ The interactions in the aromatic cage are mediated by a set of conserved polar and non-polar contacts involving amino acids at the interaction surface and by an array of water molecules present in the cavity.^2^ During our initial development of the IBPP platform, we systematically replaced amino acids with 4-azido-L-phenylalanine (AzF, **1**), a photo-crosslinkable amino acid (PCAA) in BRD4 bromodomain employing amber suppression mutagenesis.^20^ Analysis of the photo-crosslinking revealed that the L92AzF variant was remarkably capable of recognizing and crosslinking cognate binding partners upon photo-irradiation. This variant protein was employed to enrich proteome-wide interacting partners of BRD4. Given that L92 is fully conserved among BET members, we expressed corresponding L108AzF, L68AzF and L61AzF variant proteins of BRD2, BRD3 and BRDT, respectively, the closely related family members to BRD4 (Fig. 2A). Under optimized expression conditions, we observed efficient read-through of the amber codon by the evolved synthetase only in the presence of **1**. The variant proteins carrying AzF residue were confirmed by high resolution LC-MS, and purified to homogeneity and in a large quantity for binding and crosslinking experiments (Fig. 2B, S1, Table S1).

**Figure 2.**
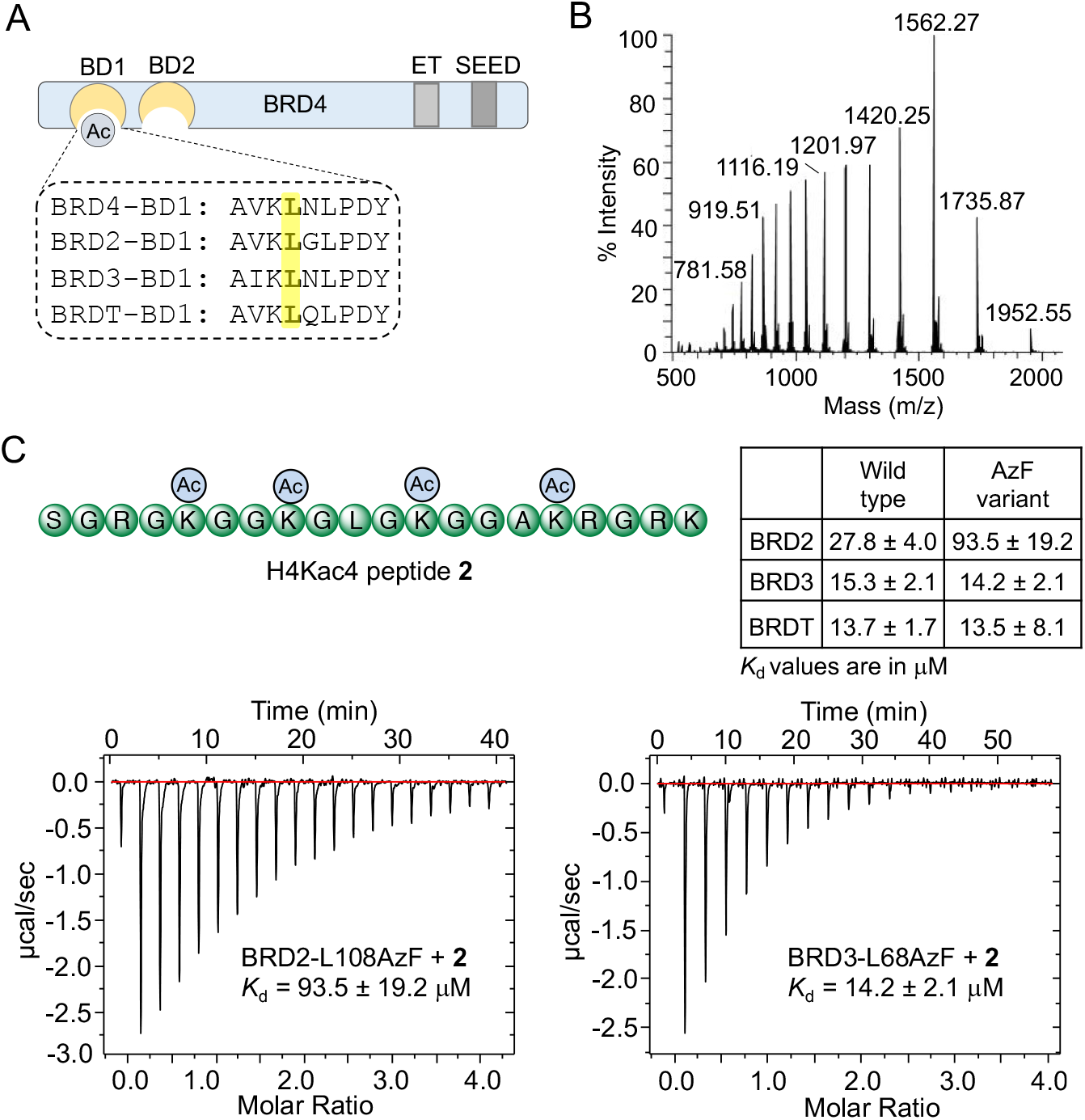
Engineering of BET bromodomains. (A) Domain structure of BET members: BRD2, 3, 4 and BRDT. L92 of the first bromodomain (BD1) of BRD4 is conserved and targeted for amber codon suppression to introduce AzF **1** to develop photoactive BRD2, BRD3 and BRDT. (B) LC-HRMS spectrum of BRD2-L108AzF mutant. See figure S1 for deconvoluted mass and the same for BRD3-L68AzF and BRDT-L61AzF variants. (C) Binding isotherms of the photoactive mutants towards H4Kac4 peptide **2** as determined by the isothermal titration calorimetry. See figure S3 for wild type BRD2, 3, BRDT and BRDT-L61AzF mutant.

To determine whether the mutants were capable of recognizing acetylated histone, we synthesized a terta acetylated histone H4 peptide (H4Kac4) **2**, corresponding to the first 21 amino acids of H4, carrying four acetyllysine residues (Fig. 2C, S2, Table S2). We determined dissociation constants (*K*_d_) of the purified proteins from H4Kac4 by isothermal titration calorimetry (ITC). The wild type BRD2, 3 and BRDT recognized the acetylated peptide with *K*_d_ of 27.8±4.0 μM, 15.3±2.1 μM and 13.7±1.7 μM, respectively, akin to previous reports (Fig. S3, Table S3).^2^ The corresponding AzF variants displayed comparable binding efficiency toward the peptide segment with *K*_d_ of 93.5±19.2 μM, 14.2±2.1 μM, and 13.5±8.1 μM, respectively (Fig. 2C, Table S3). These results demonstrate that protein engineering at a conserved site with an unnatural amino acid leads to photo-responsive bromodomain analogues for the BET members with an intact aromatic cage, while minimally altering their binding efficiency towards acetylated histone peptide.

### Crosslinking with acetylated histone peptide and full-length protein

Upon confirming the binding ability of variant proteins to their cognate acetylated partners, we examined their crosslinking potential in response to UV light. We synthesized acetylated protein segment **3** carrying a tetramethyl rhodamine (TAMRA) on the N-terminus (Fig. 3A, S2, Table S2). The wild type and photo-responsive variants of BRD2 and 3 were incubated separately with the peptide for complex formation, followed by exposure to UV light at 365 nm. Crosslinked protein bands, separated on polyacrylamide gel, were visualized using 532 nm light (*λ*_max_ for TAMRA). The variant proteins showed robust crosslinking, evident from the presence of fluorescent protein band of higher molecular weight specific to the samples exposed to UV light (Fig. 3B, S4). Under identical conditions, wild type proteins were unable to crosslink to the peptides despite their strong mutual interaction, suggesting that light-induced bond formation is indeed a result of engineering the aromatic cage with AzF in individual bromodomains. Furthermore, BET bromodomain inhibitor JQ1 at 1 μM concentration abolished the crosslinking ability of variant proteins (Fig. 3B), confirming similar binding modes for wild type and engineered proteins towards acetylated lysine residues.^21^

**Figure 3.**
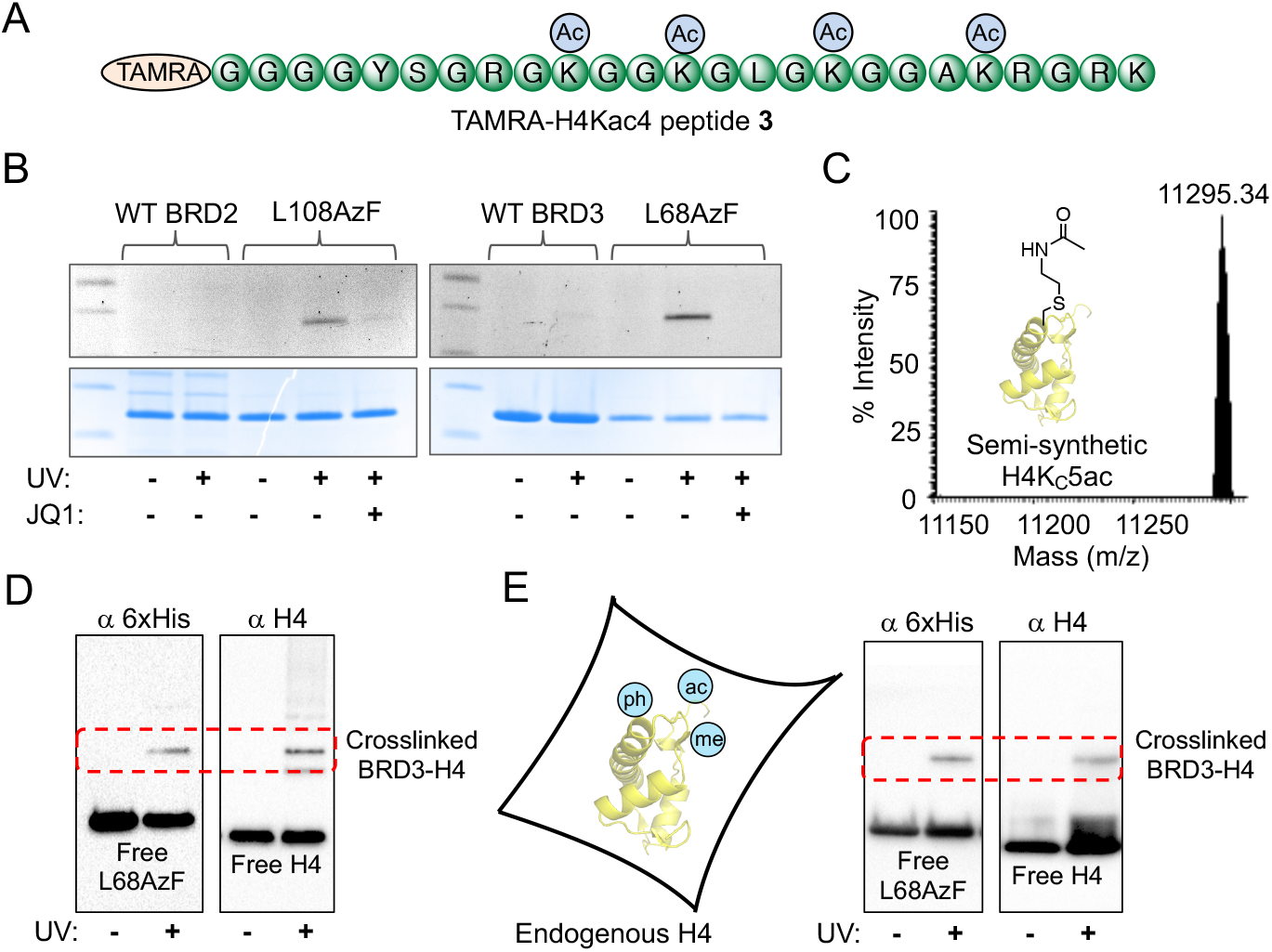
Photo-crosslinking of engineered bromodomains. (A) Sequence of TAMRA-H4Kac4 peptide **3** used for in-gel fluorescence assay. (B) Upon incubation with **3**, samples were exposed to 365 nm UV light followed by in-gel fluorescence using 532 nm light (*λ*_max_ for TAMRA). Samples that underwent successful photo-crosslinking upon exposure to 365 nm UV light are indicated by bands visible under 532 nm light. Crosslinking is significantly abolished in the presence of 1 μM JQ1. Coomassie staining of the same gel showed presence of proteins in all the samples. (C) ESI LC-MS of semisynthetic, full-length H4 carrying acetylated thialysine at position 5 (H4K_C_5ac). (D) Visualization of the crosslinking of H4K_C_5ac to BRD3-L68AzF using anti-H4 and anti-6xHis antibodies. Successful crosslinking was observed when samples were exposed to UV light. (E) Visualization of crosslinked endogenous H4 isolated from HEK293T cells to BRD3-L68AzF using anti-H4 and anti-6xHis antibodies. Successful crosslinking was observed when samples were exposed to UV light.

We further evaluated the crosslinking ability of variant proteins towards full-length H4 carrying acetylated thialysine, a lysine isostere, at position 5. We adopted a semi-synthetic method by combining protein expression and a thermal thiol-ene reaction between the H4K5C mutant and vinyl acetamide to access the histone H4 analogue.^22^ The site-specifically thia-acetylated full-length H4-K_C_5ac was confirmed by LC-MS (Fig. 3C) and subjected to photo-irradiation after incubating with BRD2 and 3 proteins. The crosslinking event was analyzed using Western blotting with both H4 and 6xHis antibodies (Fig. 3D, S4). Robust chemiluminescent signals corresponding to crosslinked species between individual variant proteins and the synthetic H4 were observed when samples were exposed to UV light.

Finally, the crosslinking ability of the BET mutants to endogenous histones was assessed. Wild-type histones were purified from human embryonic kidney (HEK) 293T cells. Such endogenous histones are biologically more relevant as they carry a full set of posttranslational modifications as opposed to its semisynthetic counterpart. The bromodomain variants L108AzF, L68AzF and L61AzF underwent efficient crosslinking with H4 present in the preparation, evident from Western blotting with relevant antibodies (Fig. 3E, S4). Taken together, these results demonstrate that upon exposure to UV, bromodomains carrying AzF at specific sites could undergo robust covalent bond formation with acetylated histone H4, as eitner a peptide segment or full-length endognous protein.

### Deciphering proteome-wide bromodomain interactome

Bromodomain analogues with site specific AzF were used to enrich novel interacting partners present in human cells. We cultured HEK293T cells in the presence of suberanilohydroxamic acid (SAHA), a broad-spectrum deacetylase inhibitor,^23^ to generate hyperacetylated human proteome. The cellular extracts were incubated with the AzF variants independently and photo-irradiated at 365 nM for 30 minutes. The crosslinked proteins were isolated using Ni-NTA beads followed by extensive washing, elution with imidazole, and resolved by SDS-PAGE (Fig. 4A). Multiple higher molecular weight protein bands were observed in Coomassie-stained polyacrylamide gels, which suggests successful crosslinking of the engineered readers to a variety of potential interacting partners present in the cellular extracts. No significant enrichment was observed in samples without exposure to UV-irradiation.

**Figure 4.**
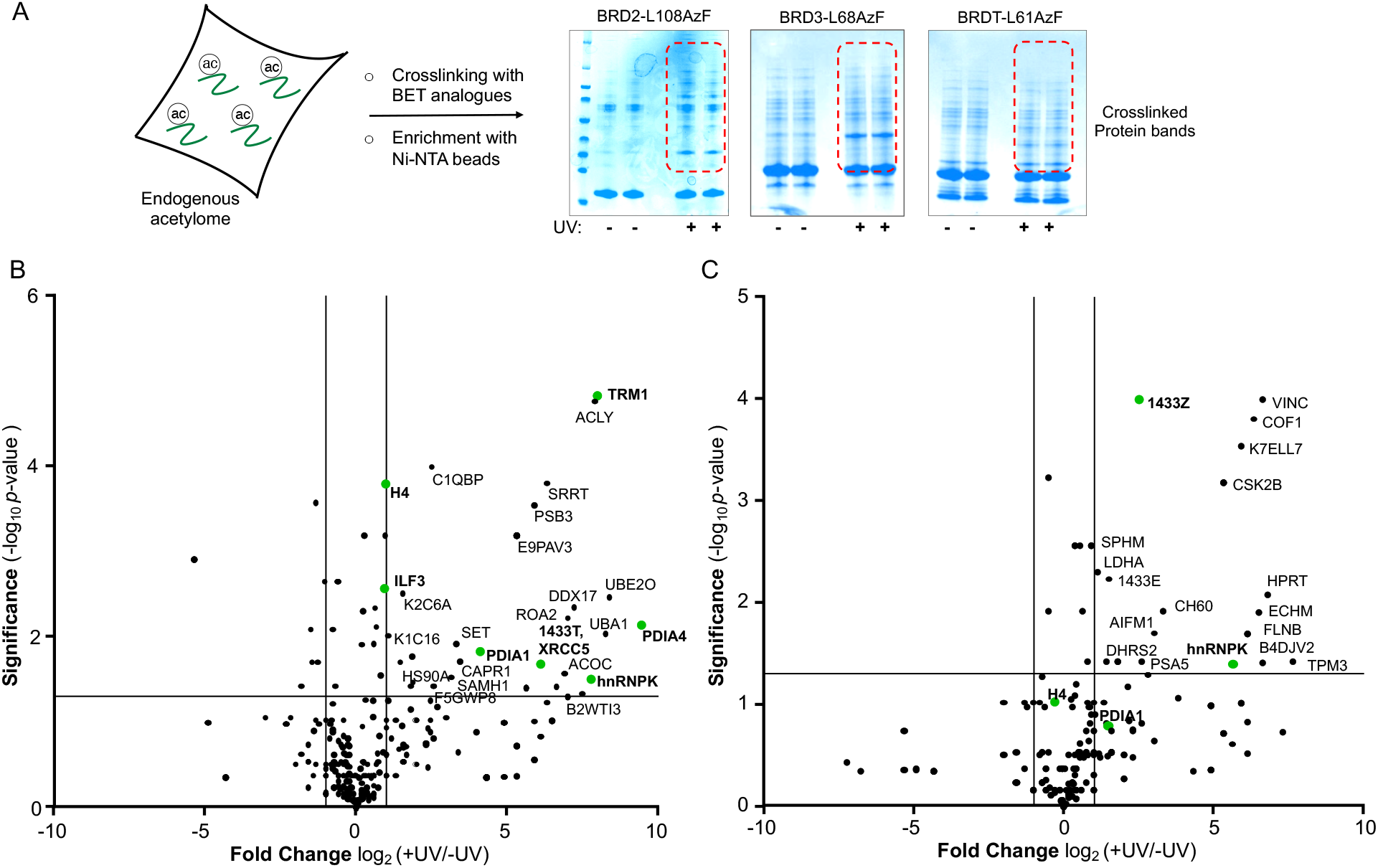
Interacting partners of BET bromodomains. (A) Schematic showing pull-down of acetylated interactome present in HEK293T cell extracts separately crosslinked with BRD2-L108AzF, BRD3-L68AzF and BRDT-L61AzF mutants. The putative interacting partners are separated on SDS-PAGE and visualized by Coomassie staining. Biological replicates are presented next to each other for both UV-treated and non-treated samples. (B, C) Volcano Plot represents potential interacting partners of BRD2 and BRD3 identified using proteomic analyses.

To characterize the putative interactome, we performed in-gel trypsin digestion of the excised gel bands. The digested peptides were subjected to liquid chromatography coupled with tandem mass spectrometry (LC-MS/MS). We performed two independent biological replicates for individual variant proteins. Duplicates of non UV-treated and identical corresponding samples were used as negative controls in parallel. We applied the following sequential criteria to narrow down the putative binding partners of the bromodomains: (i) exclusive unique peptides detected for identified proteins for individual bromodomain variant under a given condition (± light) were averaged; (ii) proteins that were present in both replicates and had a minimum average of two unique peptides were selected for subsequent analysis; (iii) the list was further narrowed by implementing a 2-fold enrichment of such candidate proteins over the negative control; and (iv) proteins within a set with a p-value of ≤0.05, obtained from a t-test between control and UV-treated samples, were considered as “high-confident” bromodomain interactome pool. Applying these criteria, we identified 68, 34, and 36 proteins as potential interacting partners of BRD2, BRD3, and BRDT, respectively, from a total of 344, 223, and 259 proteins isolated using the IBPP platform for respective bromodomains (Figure 4B, C, S5, Table S4-6). An analysis based on exclusive unique spectral counts of the enriched proteins provided similar results (Table S4-6). Furthermore, histone H4 and other known acetylated interactome of BET members such as ILF3 and PDAI1 were significantly enriched in the irradiated samples (Fig. 4B, C),^20^ validating the IBPP method for identifying authentic interacting partners of acetyllysine readers from relevant cell extracts.

Using the IBPP approach, we identified both nuclear and cytosolic proteins as putative interacting partners of BRD2, 3 and BRDT. Analysis of the exemplary BET interactome with high scores (p-value of ≤0.05) revealed that the bromodomain partners spread across a broad spectrum of signaling pathways (Fig. 4, Table S4-6). The interactomes include DNA-binding proteins such as replication factors (EF2, EF1D, FUBP2/3), transcription factors (BCLAF1, FUBP2, ILF3), histones (H2A, H4), and DNA repair proteins (XRCC5); RNA-binding proteins such as RNA methyltransferases (TRM1), splicing factors and helicases (DHX15, DDX17, SF3B3), polyA tail-binding proteins (PABP1) and ribonucleoproteins (hnRNPK); metabolic enzymes (SHMT, ACLY); foldase and isomerase (HSP60, PDIA1); ubiquitin activating and conjugating enzymes (UBA1, UBE2O, UBTL3); and scaffolding and structural proteins (14-3-3T, cofilin). These results suggest that the newly deciphered interactome orchestrates a diverse set of cellular processes including DNA replication, transcription, DNA repair, RNA splicing and stability, cytoskeleton dynamics, ubiquitin proteasome pathway, cellular homeostasis and apoptosis, and oxidative and metabolic processes. Hence, each interactome reported here expands the canonical functions of BET members far beyond chromatin and transcription. We, however, do not claim the presented catalog to be an exhaustive registry of interacting partners of BRD2, 3 and BRDT, as the IBPP-derived interactome covers some, but not all, of the known interacting partners such as GATA1 and POLR2A. We surmise that the inconsistencies may arise from differential composition of certain partners in HEK293T cells and/or executed experimental strategies involving specific deacetylase inhibitors, which can lead to distinct ‘acetylome’ patterns in cultured human cells.^24^

Analysis of the interactomes revealed a significant overlap between the interacting partners of BET members, consistent with their sequence and structural similarities. Proteins such as PDIA1 and 4, hnRNPK, ILF3, members of 14-3-3 family, and proteins in proteasome and ubiquitin pathways are common to the BRD2-3 and BRDT bromodomains. Our previous work documented that the interactome of BRD4 includes some of these partners, particularly ILF3, hnRNPK, PDIA1 and PDIA4, indicating potential overlapping functions of the BET bromodomains via shared non-histone interactomes. The current study also led to distinctive protein-protein interactions mediated by the BET members. For example, while BRD2 interacts specifically with DNA-binding proteins FUBP2 and EF1D, BRD3 engages with the apoptosis factor AIFM1 and cytoskeletal protein cofilin, and BRDT recognizes splicing factor SF3B3 and Bcl2-associated transcription factor 1 (BCLAF1). Our findings highlight that bromodomains could potentially regulate distinctive pathways by binding to unique acetylated proteome.

### Biochemical evaluation of the acetylated interactome

Recent unbiased proteomic studies have identified several thousands of proteins to be acetylated at specific lysine residues in human cells.^24^ However, the nature of the writers (acetyltransferases) and readers (bromodomains) dedicated towards marking and recognizing the acetylome (collection of acetylated proteins) is largely unknown. Scrutiny of the acetylome in available databases along with a survey of primary literature revealed that majority of the proteins identified by the IBPP platform are indeed targets of acetylation, often at multiple lysine residues, in human and other mammalian cells. We sought to determine if BET bromodomains could recognize acetylated lysine residues in these proteins with *in vitro* binding assays. We selected ten proteins identified with high scores in our proteomic experiments and synthesized the corresponding short peptides **4**-**13**, each carrying at least one acetyllysine at known acetylated sites in human cells (Fig. 5A, S6, Table S2). These protein segments were chosen with varied acetylation patterns, both canonical histone-like and non-canonical motifs, to gain insights to the sequence specificity and mode of interaction by the BET bromodomains.

**Figure 5.**
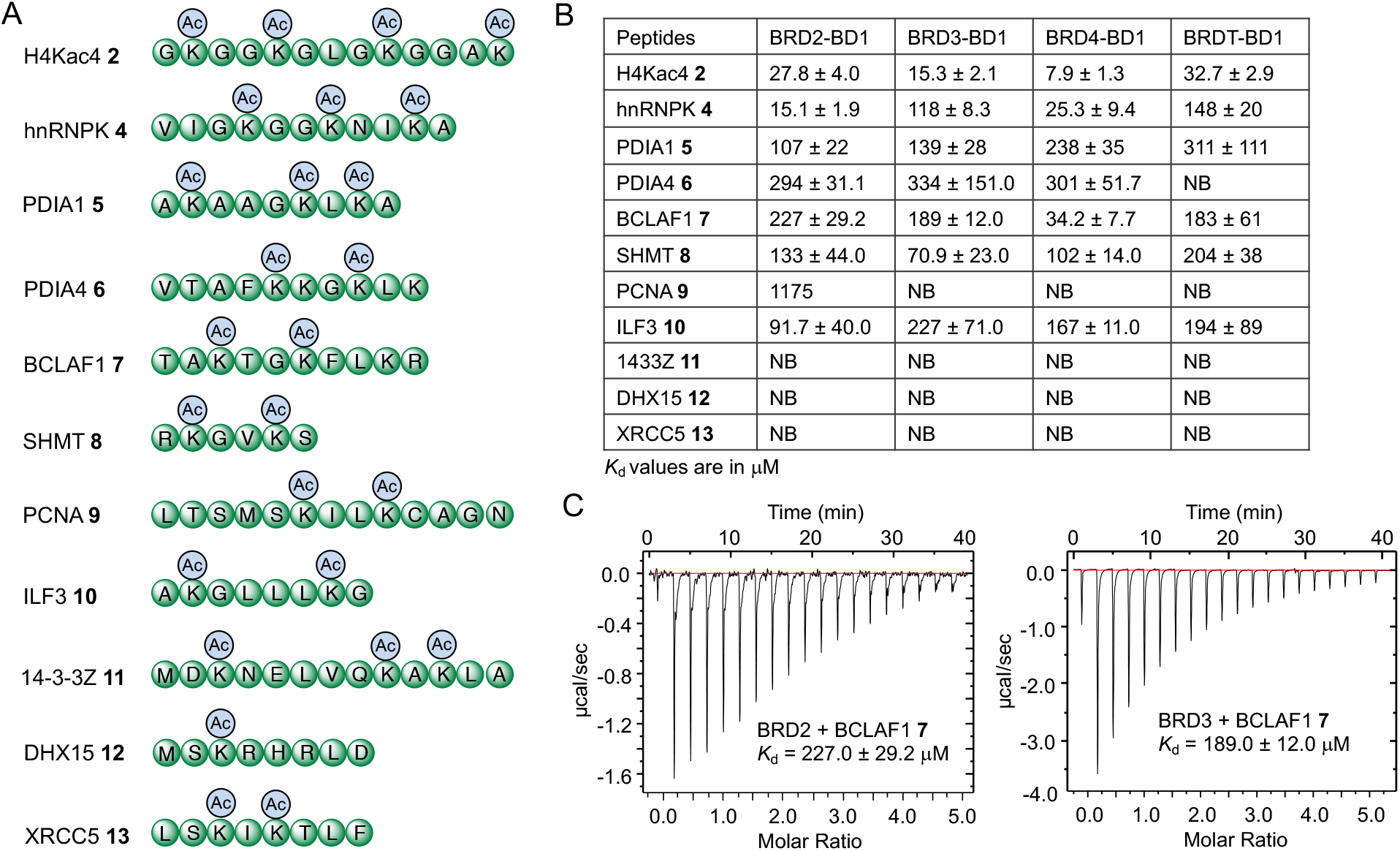
Biochemical assessment of novel BET interactome. (A) Sequences of synthesized protein segments carrying one or more acetyllysine units. See Figure S6 for corresponding MALDI-MS spectra. (B) Solution phase dissociation constants of peptides from first bromodomains (BD1) of BRD2-4 and BRDT were measured using isothermal titration calorimetry (ITC) as shown in the table in (B). See Figure S7 for the individual binding isotherm. (C) Representative binding isotherms of the BCLAF1 peptide **7** towards BRD2-BD1 and BRD3-BD1, respectively, as determined by ITC. (NB = No binding)

To obtain accurate binding information, we determined dissociation constants (*K*_d_) for the first bromodomain of BRD2-3, BRDT and the set of selected acetylated protein segments by isothermal titration calorimetry (ITC). We also included the first bromodomain of BRD4 to examine interactome promiscuity and specificity among the highly conserved BET family members. All four of these proteins recognized the cognate histone peptide H4ac4 **2** strongly with *K*_d_ closer to the reported values, confirming their biochemical integrity (Fig. 5B, S7, Table S3). However, the bromodomains displayed varied strengths of interaction towards the acetylated non-histone peptides, as determined by ITC. Short segments carrying H4-like diacetyl motifs (e.g. K5ac<u>GG</u>K8ac, K8ac<u>GLG</u>K12ac and K12ac<u>GGA</u>K16ac) displayed strong interactions in the binding assay. Significant binding to the readers was observed for hnRNPK (Kac<u>GG</u>Kac) **4**, PDIA1 (Kac<u>AAG</u>Kac) **5**, PDIA4 (Kac<u>KG</u>Kac) **6**, and BCLAF1 (Kac<u>TG</u>Kac) **7** segments, with BRDT displaying a slightly lower affinity compared to other BET members (Fig. 5B–C, S7, Table S3). In our panel, we also included a short peptide **8** carrying Kac<u>GV</u>Kac motif from SHMT, previously shown to be recognized by BRD4 bromodomain. The peptide was indeed recognized by the set of BET proteins with comparable affinity. We also synthesized non-acetylated protein segments corresponding to ILF3 and BCLAF1 as representative examples, and observed no binding to the wild type BRD4; the binding data corroborates the notion that interactome recognition in BETs relies on acetylated lysine residues in their partners.

We next turned our focus to the bromodomain interacting partners that do not harbor a canonical H4 motif. Among these unique sequences, ILF3 peptide **10** with a spacer of four amino acids, AKac<u>GLLL</u>KacG, displayed strong binding to the four selected bromodomains (Fig. 5B, S7, Table S3). In contrast, the acetylated peptides derived from 14-3-3Z **11**, DHX15 **12**, and XRCC5 **13** failed to bind the bromodomains. A closer look at their sequences revealed that 14-3-3Z peptide **11** carries multiple acidic amino acids that could bar binding by posing electrostatic repulsion with the conserved aspartic acids (e.g., D72, D120, D121 in BRD3 and D96, D144, D145 in BRD4) that surround the aromatic cage in bromodomains, and interact with basic histone proteins. DHX15 segment **11** contains a single acetyllysine, and the two acetyllysines in XRCC5 **13** are separated by a single amino acid, rather than by the typical 2-3 amino acid linker present in H4. We posit that these anomalies in DHX15 and XRCC5 primary sequences are potential reasons for their compromised affinity towards the readers. In summation, by examining only a set of ten protein segments, we have biochemically confirmed six unique sequences as potential interacting partners of BRD2-4 and BRDT. Such a score indicates that additional BET interactome within the panel of high confidence proteins identified using IBPP platform will be identified by examining defined peptide sequences carrying acetyllysine units. More importantly, some of the validated protein segments such as ILF3 do not contain the canonical Kac<u>GG</u>Kac histone pattern, thus expanding the sequence repertoire for engaging bromodomains. Such unique interactions are likely to dictate the biological functions of BET bromodomains beyond chromatin-dependent processes.

### Structural insights into novel acetyllysine-bromodomain interactions

To gain insight into the atomic interactions between BET bromodomains and the newly identified interactome, we sought to determine crystal structures of acetylated protein segments bound with the readers. We focused on the first bromodomains of BRD3 and 4 as representative BET members. To obtain crystals of the complexes, we purified about 20 milligrams of recombinantly expressed tag-less individual bromodomains. Short segments, corresponding to hnRNPK, SHMT, ILF3 and BCLAF1, encompassing diverse sequence motifs and biological functions, ranging from transcription to cell division and cellular metabolism, were selected for structural studies. Our efforts led to a total of eight high-resolution crystal structures of BRD3 and 4 bound to these acetylated non-histone segments. A summary of the crystallization conditions, data collection, refinement statistics and RSCB accession codes is provided in Table S7.

Analysis of the newly determined crystal structures revealed several conserved features among the bromodomain family. These include the presence of a left-handed, four-helix bundle termed the ‘bromodomain fold’, consisting of four α helices (αZ, αA, αB, and αC) (Fig. 6–8, S8); the two interhelical loops of variable length, known as the ZA and BC loops, connect the αZ and αA, and αB and αC helices, respectively, to form a hydrophobic pocket, the ‘aromatic cage’, which accommodates acetylated lysine residues. Within this deep central cavity, the carbonyl oxygen of the acetyl group on the first Kac residue in the newly identified partners forms a critical hydrogen bond with the sidechain amide nitrogen (N_δ2_) of a highly conserved asparagine residue (N140 in BRD4 and N116 in BRD3; Fig. 6–8 and S8) in the bromodomain family. The structures also revealed an array of water molecules in the acetyllysine binding pocket, where they form an extensive network of hydrogen bonds to mediate interactions between acetyllysine and the reader.

**Figure 6.**
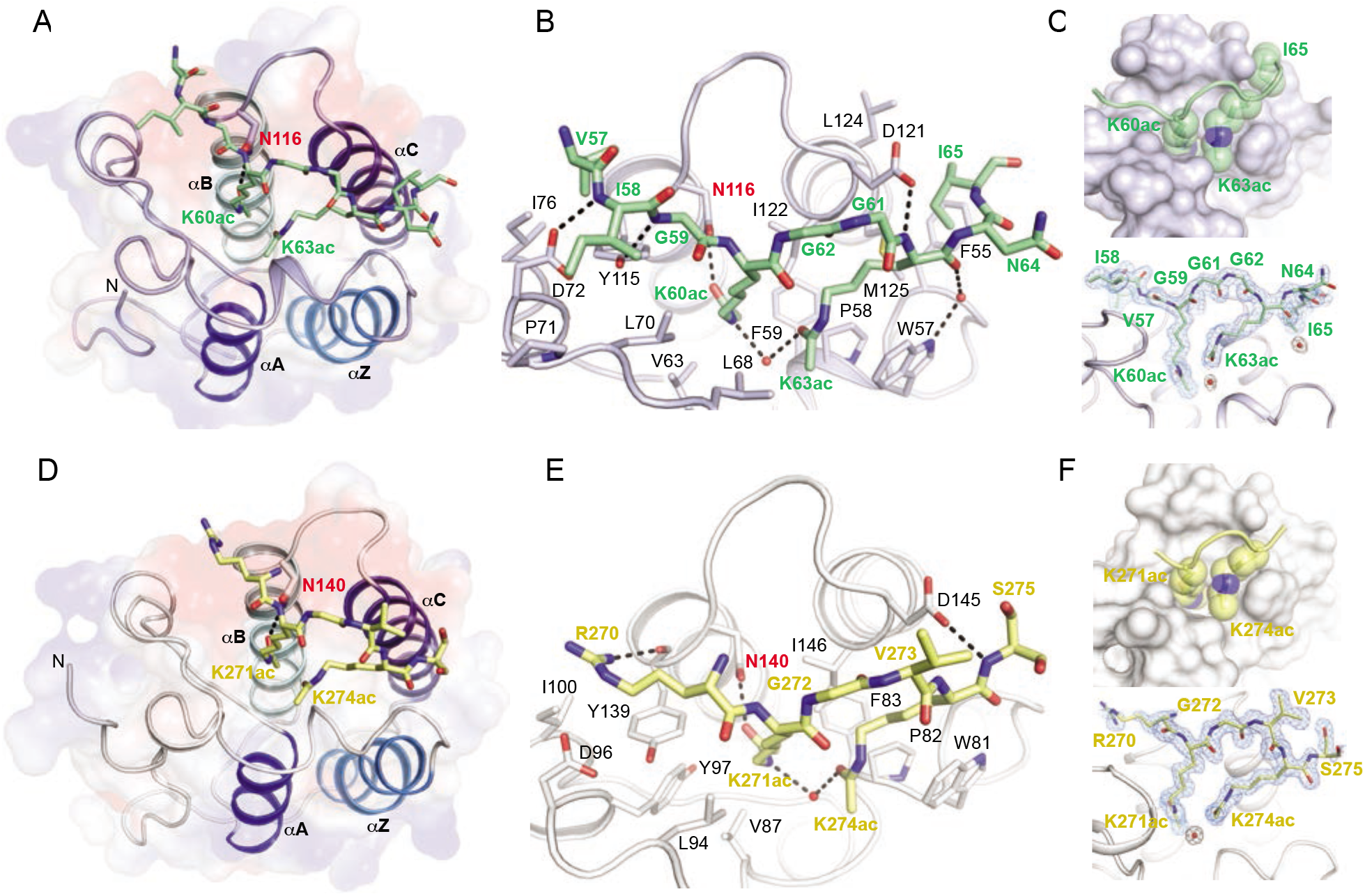
Structures of BRD3-hnRNPK and BRD4-SHMT complexes. (A-C) Closeup view of the BRD3-BD1 (light blue) complxed with hnRNPK (green) (PDB: 7RJK). A continuous 9 amino acid long acetylated (at Lys60 and Lys63) hnRNPK peptide that interacts with side chains of BRD3 are shown in stick representation. Surface representation depicts binding of K60ac and K63ac sidechains of hnRNPK into the aromatic cage of BRD3. Electron density from simulated annealing composite omit (5% omission of the model) maps contoured at 1.2 σ (blue mesh) are shown for the bound acetylated protein segment. (D-F) A view of BRD4-BD1-SHMT complex containing a continuous 6 amino acid long acetylated (at Lys271 and Lys274) SHMT segment (yellow) (PDB: 7RJP). Interacting side chains in both proteins are depicted in stick representation. Surface representation highlights binding of K271ac and K274ac sidechains of SHMT into the aromatic cages of BRD4. Composite omit maps for the bound acelytaed SHMT is shown in blue mesh. Potential hydrogen bonds are denoted with black dashed lines. All figures depicting structure were generated with PyMol.^25^ See figure S8 for high-resolution strcutures of BRD3-SHMT (PDB: 7RJL) and BRD4-hnRNPK (PDB: 7RJO) complexes.

In addition, the structures also revealed a noticeable difference in the mode of binding for each peptide. In the complexes of hnRNPK and SHMT, carrying KacGGKac (aa 60-63) and KacGVKac (aa 271-274) motifs, respectively, the acetyllysine pairs are inserted into the aromatic cage of bromodomains (Fig. 6, S8). The mutual interactions recapitulate those observed in H4-bromodomain (PDB ID 3UW9), including the one with conserved asparagine. We surmise that the similarities in binding between hnRNPK/ SHMT and H4 are driven by sequence similarities within the respective motifs. Thus, our analyses strongly suggest that BET bromodomains are capable of recognizing Kac-XX-Kac motifs outside histone H4. The hnRNPK and SHMT complexes, however, reveal distinct sets of interactions: In SHMT, R270 preceding the first Kac (K271) interacts with conserved D96 in BRD4 bromodomain (D72 in BRD3) (Fig. 6E, S8), whereas isoleucine (I64), following second Kac in hnRNPK, inserts into the extended aromatic cage and engages in hydrophobic interaction with conserved residues (F79, L148 and M149 in BRD4) in bromodomains, thus sterically filling in the residual volume of the cavity (Fig. 6B, S8).

In the segment sequence of ILF3, two acetyllysine residues are separated by four amino acids: KacGLLLKac (aa 100-105). This raises the question of how bromodomains interact with this sequence motif, which is distinct from the canonical histone H4. The structure of BRD3-ILF3 complex reveals that the first Kac of ILF3 segment makes a typical contact with conserved asparagine N116 and adopts a H4-like spatial configuration using an analogous ‘GL’ motif (Fig. 7A, B). This arrangement allows the second leucine of ILF3 peptide, L103, to enter the aromatic cage where K8ac of H4 resides in canonical BRD3-H4 complex (Fig. 7C). The third leucine (L104) in the peptide is packed in the extended cavity comprising of W57, F59 and M125. No appreciable electron density was observed for the second acetyllysine (K105ac) in the BRD3-ILF3 complex. In the structure of the BRD4 complex, however, the interaction of ILF3 is noticeably different (Fig. 7D, E). Although contact between the conserved N140 in aromatic cage of BRD4 and the first acetyllysine of ILF3, K100ac, is maintained, neither L103 nor L104 are engaged at the interface; instead, the leucine “duo” adopts a short turn, which allows the first leucine, L102, and the second acetyllysine, K105ac, of ILF3 to pack together into the aromatic cage (Fig. 7F). It is important to highlight that the two acetyllysines of ILF3 engage in a water mediated interaction similar to K5ac-K8ac interaction observed at the BRD4-H4 interface (Fig. 7E). The structural analyses exemplify how bromodomains exploit diverse and distinct binding mechanisms to interact with an array of distinct non-histone proteins.

**Figure 7.**
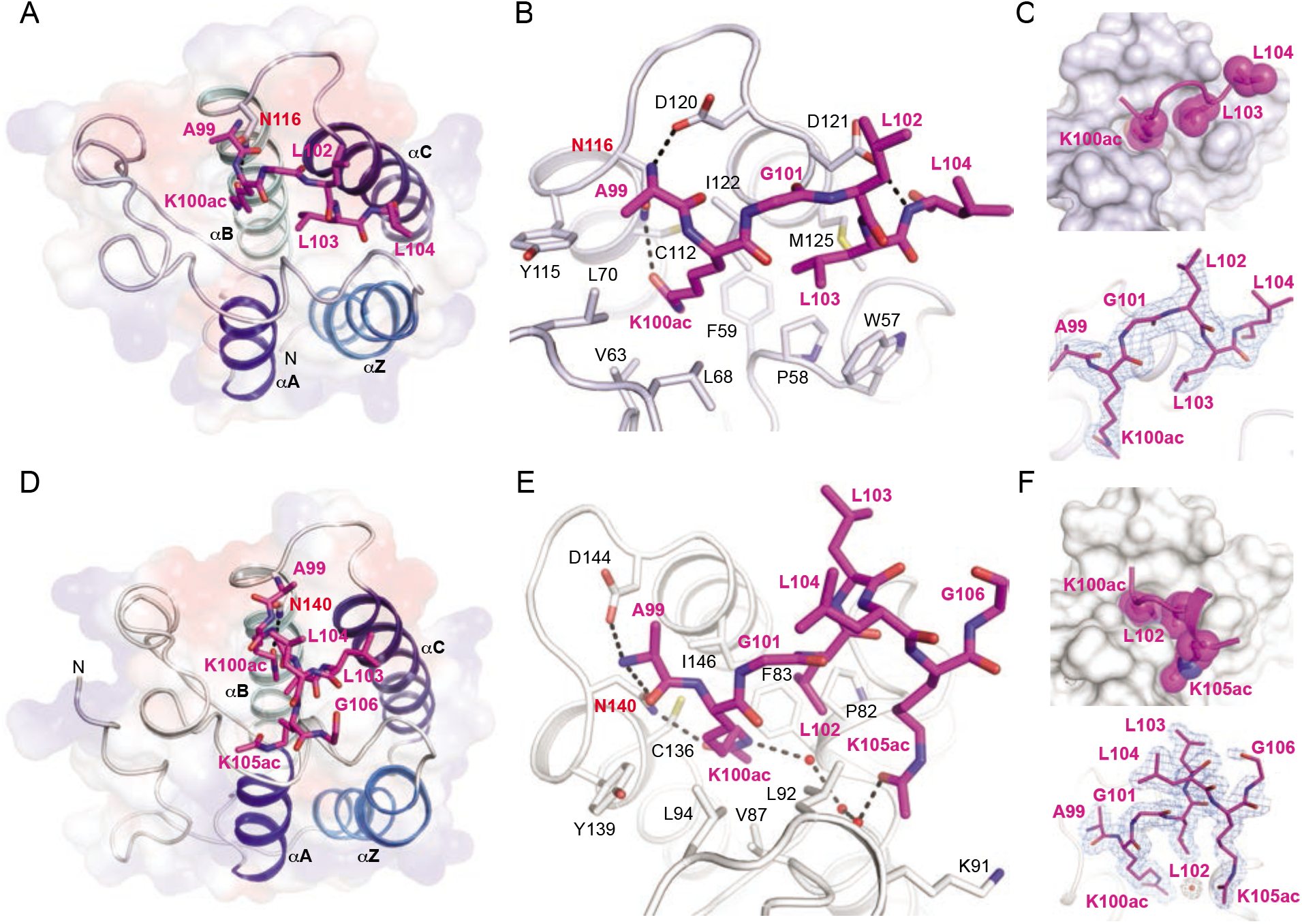
Structures of BRD3 and 4 in complex with acetylated ILF3. (A-C) A view of BRD3-BD1 (light blue) complexed with ILF3 peptide (in pink) with acetylated Lys100 (PDB: 7RJM). The interacting ILF3 segment and BRD3 side chains are shown in stick representation. Surface representation and electron density (simulated annealing composite omit) maps show binding of K100ac and L103 sidechains of ILF3 into the aromatic cage of BRD3. (D-F) Interaction of BRD4-BD1 with ILF3 (PDB: 7RJQ). A seven amino acid long acetylated (at Lys100 and Lys105) segment of ILF3 (in pink) and side chains of BRD4 that interact with the segment are shown. Surface representation and composite omit maps show binding of K100ac, L102 and K105ac sidechains of ILF3 into the aromatic cages of BRD4.

The BCLAF1 peptide segment encompasses the KacTGKac (aa 332-335) amino acid array, which closely resembles the canonical KacGGKac motif of histone H4. However, surprisingly and rather remarkably, the bromodomains employed a distinct recognition mode to bind to BCLAF1 (Fig. 8). The structures highlight that the second acetyllysine, K335ac, and not the first acetyllysine, K332ac, of BCLAF1 segment contacts the conserved asparagine (N116 in BRD3 and N140 in BRD4) in the cage, while the adjacent phenylalanine (F336 in BCLAF1) is inserted into the space typically occupied by the second acetyllysine, K8ac, of H4 (Fig. 8B, E). No appreciable electron density was observed for the first acetyllysine, K332ac, in either of the BCLAF1 complexes, suggesting the BET-BCLAF1 interactions are driven by a KacØ motif (where Ø is a bulky hydrophobic residue). The unique binding mode induces a short turn in the BCLAF1 segment accompanied by structural changes on the surface of bromodomains typically not observed in the interactions with histones (Fig. 8C, F). It is, however, noteworthy to state that the binding mode is somewhat similar to the interaction of K221acY222 motif of BAZ1b (PDB ID 4NNF) with BRD4, suggesting such noncanonical recognition pattern is widespread among bromodomain family members.^3^

**Figure 8.**
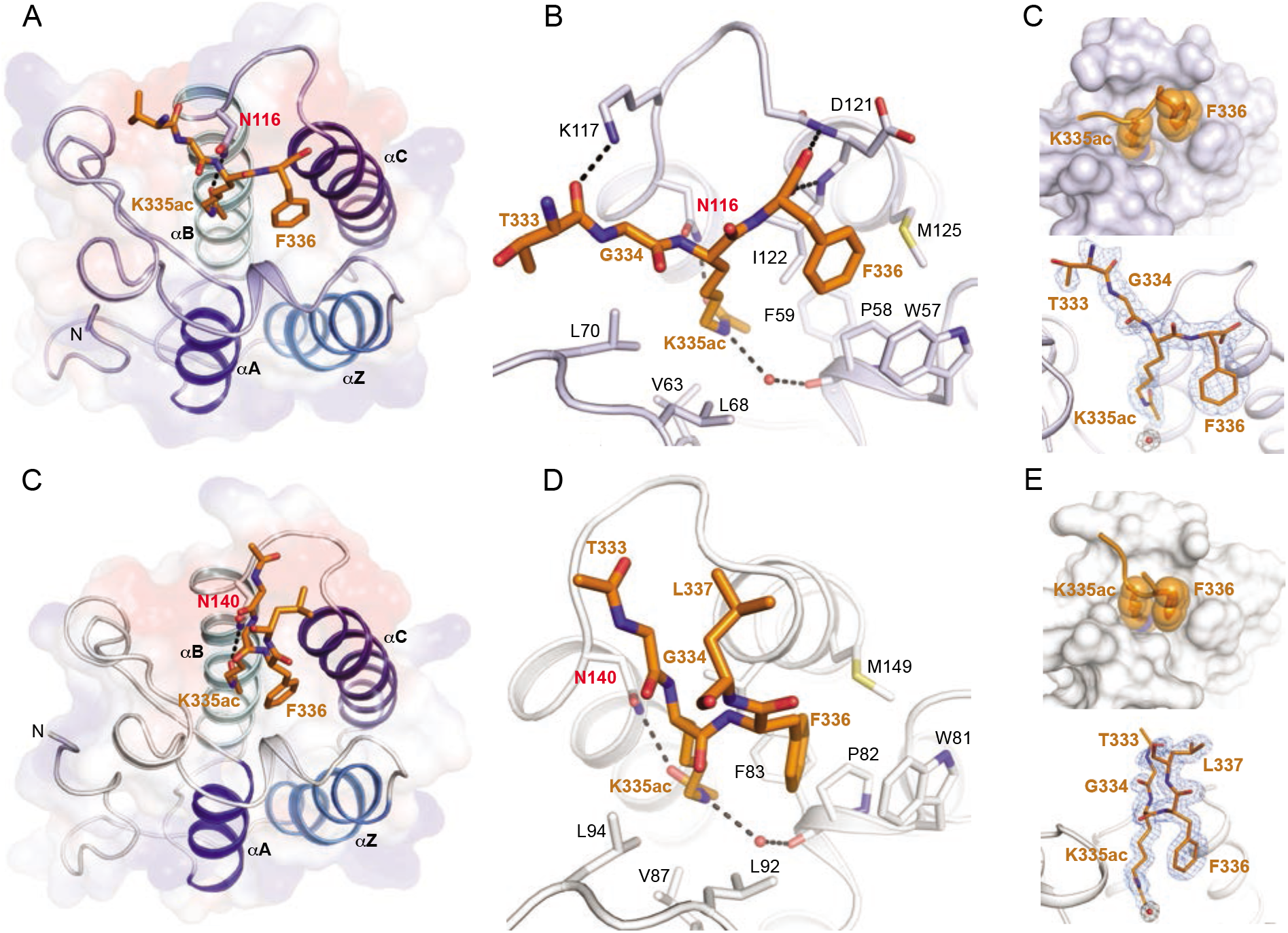
Structures of BRD3 and 4 in complex with acetylated BCLAF1. (A-C) A view of the BCLAF1 interactions with BRD3-BD1 (PDB: 7RJN). The acetylated BCLAF1 (at Lys335) segment (4 amino acid long; colored gold) and BRD3-BD1 side chains are shown in stick representation. Surface representation and composite omit maps show binding of K335ac and F336 sidechains of BCLAF1 into the aromatic cage of BRD3. (D-F) A closeup view of the BRD4-BD1 and BCLAF1 interactions (PDB: 7RJR). The acetylated BCLAF1 (at Lys335) segment (4 amino acid long; colored gold) interacting with BRD4 side chains are shown in stick representation and labeled. Surface representation and electron density maps (simulated annealing composite omit) highlights binding of K335ac and F336 sidechains of BCLAF1 into the aromatic cage of BRD4.

Taken together, the structural data presented here strengthen our proteomic and biochemical findings on non-histone interactome of BET bromodomains and uncovered novel binding modes within the interactome. More importantly, our results demonstrate that associations of BET bromodomains are mediated via diverse protein sequences and structural determinants besides those residing in histones. Perhaps the most striking structural observations were the interactions between ILF3 and BCLAF1 segments in which hydrophobic sidechains substituted acetyllysines of the canonical KacGGKac motif. It is tempting to speculate that non-histone partners without an acetyllysine but with complementary hydrophobicity could dock into the aromatic cages of BET bromodomains and expand the BET interactome repertoire even further. We surmise that the existence of potential non-acetyllysine binding arrays (ØX[?]Ø) are perhaps more prevalent for orphan bromodomains whose interacting partners are yet to be identified.

### Analysis of ILF3-BRD4 interaction in cultured human cells

We next sought to investigate the mutual interaction between ILF3 and BRD4 along with its potential role in establishing novel signaling pathways to regulate downstream processes in cells. ILF3 and BRD4 share several common features: chromosomal location, expression pattern, nuclear localization, euchromatin association, G2/M transition, and role as host factors for regulating viral gene replication.^26–30^ However, a direct link between the two gene products in controlling cellular processes has not been established. We focused on nuclear factor 90 (NF90) and 110 (NF110) isoforms of ILF3, which harbor a range of protein-protein interaction domains.^31^ NF110 is further characterized by additional RNA-binding domains at the C-terminal. It has been shown that K100 in the DNA zinc finger domain (DZF) of NF90/110 undergoes robust acetylation in human cells, but the cellular role of this posttranslational modification is undefined.^24^ Prompted by our biochemical and structural results demonstrating the unique binding between the BD1 of BRD4 and KacGLLLKac (aa 100-105) of ILF3, we expressed Flag-tagged ILF3 in SAHA-treated HEK293T cells via transient transfection. Immunoprecipitation (IP) with anti-BRD4 antibody followed by Western blot using Kac antibody, revealed that ILF3 indeed undergoes lysine acetylation in cells (Fig. 9A). Acetylation was detected only in the presence of exogenously expressed acetyltransferases GCN5 and CBP, unleashing the nature of the potential acetyllysine writers in ILF3. More importantly, IP experiment with an ILF3 variant carrying alanine mutation at K100 significantly reduced acetylation level, further confirming K100 as the potential sites of acetylation, consistent with reported observations (Fig. 9A).^24^

**Figure 9.**
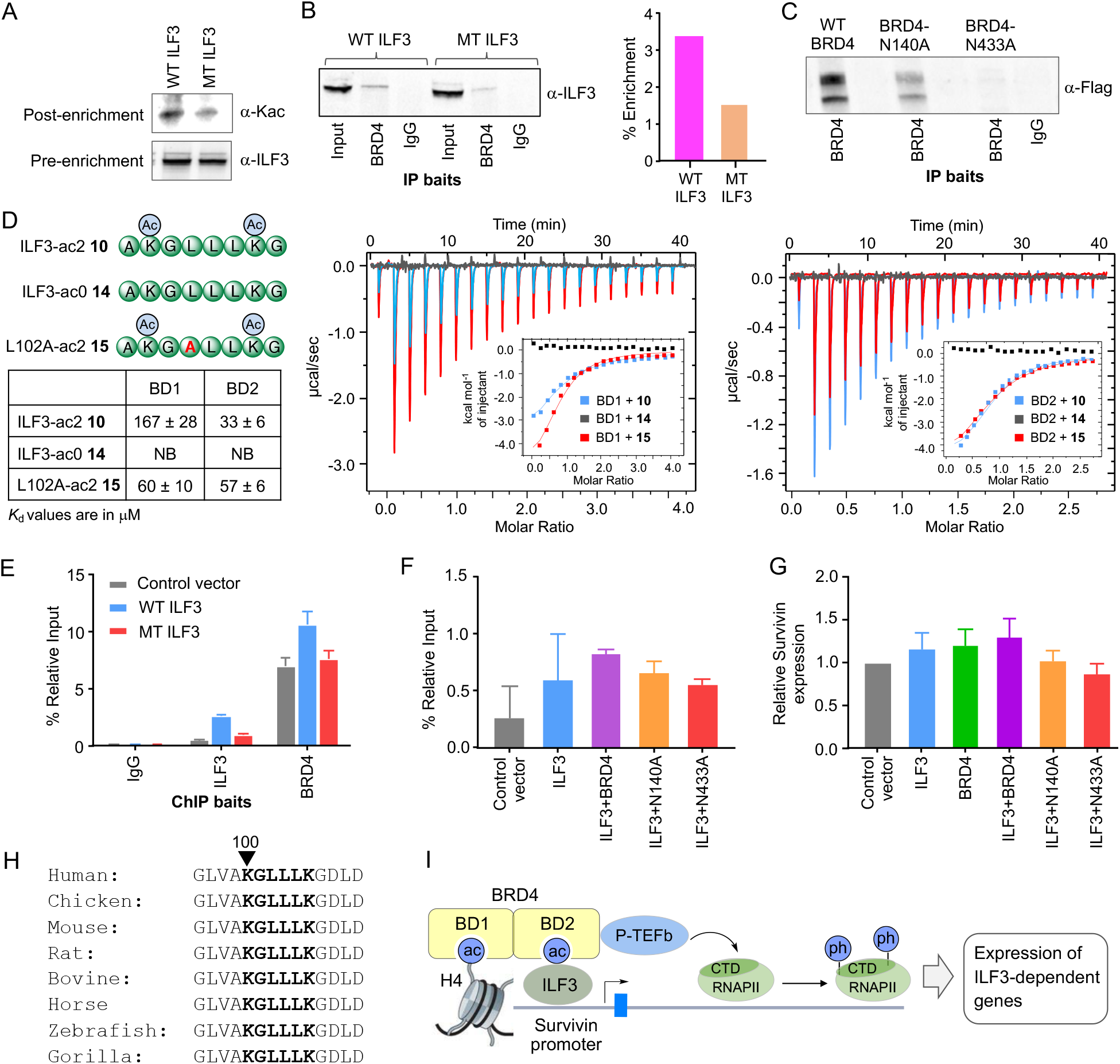
Validation of BRD4-ILF3 interaction in human cells. (A) Immunoprecipitation with BRD4 antibody followed by Western blot using Kac antibody shows ILF3 acetylation at Lys100. Acetalytion level decreases with K100A mutation of ILF3. (B) Immunoprecipitation with BRD4 antibody followed by Western blot using ILF3 antibody confirms ILF3-BRD4 interaction. K100A mutation significantly decreases the association. (C) Immunoprecipitation with BRD4 antibody followed by Western blot using Flag antibody shows ILF3-BRD4 interaction decreases with defunct BRD4 mutations. (D) Dissociation constants of diacetylated ILF3 peptides (wild type **10** or L102A variant **16**) from the first (BD1) and second (BD2) bromodomains of BRD4 as judged by isothermal titration calorimetry. (E-F) Chromatin immunoprecipitation (ChIP) followed by qPCR showing localization of ILF3 and BRD4 to Survivin promoter. Recruitment of ILF3 to Survivin locus decreases with defective BRD4 mutations. (G) ILF3-BRD4 interaction synergistically regulates Survivin expression as revealed by qRT-PCR. BRD4 mutations decrease the expression of Survivin. (H) Sequence conservancy of ILF3 KGLLLK (100-105) among representative eukaryotes. (I) Proposed model for multivalent interaction of BRD4 with acetylated histone and ILF3 via BD1 and BD2, respectively, for locus-specific recruitment leading to transcriptional activation of ILF3 targets through P-TEFb (CDK9)-mediated phosphorylation of RNAPII.

To examine if the acetylated ILF3 is capable of interacting with BRD4 in cells, we performed IP experiments with cells overexpressing ILF3. We indeed detected ILF3 in Western blots when enriched using BRD4 antibody (Fig. 9B). Similarly, in reversed pull-down using the ILF3 antibody, endogenous BRD4 was enriched as evident from Western blot analysis (Fig. S9). IP with immunoglobulin G (IgG) failed to enrich either protein, further confirming the mutual interaction between ILF3 and BRD4. To demonstrate that atypical interaction is indeed mediated by the KacGL (aa 100-102) motif as revealed by our structural analysis, we generated a K100A/L102A double mutant of ILF3. In subsequent immunoprecipitation experiments, the double mutant failed to interact with endogenous BRD4 as the signal for ILF3 diminished significantly in Western blot when enriched using BRD4 antibody (Fig. 9B). The lower enrichment of the mutant ILF3 was neither due to reduced expression of the ILF3 as shown by qRT-PCR using ILF3-specific primers, nor was it because of any alteration in antibody recognition as discernable from the Western blot analysis (Fig. S10). Taken together, our results demonstrate that the two transcriptional regulators interact in human cells and their mutual interaction critically relies on the acetylated KGLLLK (aa 100-105) motif in the DZF domain of ILF3.

After confirming BRD4-ILF3 interaction in cells and the potential site of association on ILF3, we sought to gain further insights into the binding mechanism of full-length BRD4 carrying two bromodomains. N140 and N433 are two residues in the first (BD1) and second (BD2) bromodomains, respectively, that are critical for acetyllysine recognition. We generated the N140A and N433A independently in full-length BRD4 and expressed individually in HEK293T cells. Subsequent IP using BRD4 antibody followed by Western blot for ILF3 revealed that both mutations decreased the interaction between the transcription factor compared to wild type BRD4 (Fig. 9C). Remarkably, the N433A mutation had a more pronounced effect on binding, suggesting a dominant role of BD2 in interacting with ILF3 in cells. To confirm the in-cellulo binding mode, we measured dissociation constants of ILF3 peptide **10** from bacterially expressed BD2. As expected, BD2 failed to bind the non-acetylated ILF3 **14**, but showed robust binding towards the diacetylated peptide **10** with a *K*_d_ of 33 ± 6 μM (Fig. 9D, Table S3). Remarkably, the affinity is significantly stronger than that of BD1 (*K*_d_ = 167 ± 28 μM) (Fig. 5B), corroborating well with the *in cellulo* results. To further probe the binding difference between BD1 and BD2 towards ILF3, we synthesized an ILF3 variant peptide AKacG<u>A</u>LLKacG (aa 99-106) **15** carrying L102A mutation (Fig. 9D). To our surprise, this mutation led to stronger binding of the peptide to BD1 (*K*_d_ = 60 ± 10 μM) and a reduced affinity toward BD2 (*K*_d_ = 57 ± 6 μM), suggesting a role for L102 in domain specificity (Fig. 9D, Table S3). Intriguingly, the L102A mutation makes the ILF3 sequence closer to H4 (GKGGAK), thus enhancing its interaction with BD1. Our biochemical results further support a prevailing model in which BD1 interacts with the acetylated histones, whereas BD2 engages with the non-histone proteins for multivalent BRD4-chromatin association to regulate downstream processes (Fig. 9I).^10^

### ILF3-BRD4 interaction regulates transcription of mitotic gene

Upon validating the ILF3-BRD4 interaction in cells and elucidating their binding mechanism, we intended to determine the function of this novel association, particularly in the context of transcriptional regulation. Both ILF3 and BRD4 are required for mitotic progression and cytokinesis, and play critical role in mediating G2/M transition. Survivin is an evolutionarily conserved eukaryotic protein that is essential for cell division, ensuring proper attachment of microtubule filaments to kinetochore for sister chromatid separation and mitosis.^32,33^ Its expression peaks at G/M phase and is shown to be independently regulated by ILF3 and BRD4.^32,34–36^ Although a combined role of BRD4 and NF-κB in Survivin expression is known, it is not clear how ILF3 regulates Survivin at the transcriptional level. We sought to examine if the ILF3-BRD4 duo is mechanistically linked to Survivin expression. We performed chromatin immunoprecipitation (ChIP) with IgG, ILF3, and BRD4 antibodies from cells expressing wild type or binding deficient (K100A/L102A) NF90/ILF3, followed by qPCR using primers corresponding to the Survivin promoter. The endogenous ILF3 and BRD4 indeed localized at the Survivin promoter, as revealed by higher enrichment of the promoter compared to IgG control (Fig. 9E). Furthermore, overexpression of wild type ILF3 led to enhanced localization of both proteins; in contrast, the ILF3 mutation reduced their occupancy to the locus, confirming that acetyllysine-mediated mutual ILF3-BRD4 interaction is critically important to their recruitment at the Survivin locus (Fig. 9E). BRD4 has been shown to have an alternative mechanism, such as interaction with p65 component of NF-κB complex, for localization to the Survivin locus,^35^ which partially compensates for the loss of its interaction with ILF3. To gain further mechanistic insights into the promoter localization of ILF3-BRD4, we performed ChIP from cells expressing either wild type or mutant BRD4 (N140A or N433A) using ILF3 antibody. While the wild type reader enhanced localization of NF110 to Survivin promoter as expected, the N433A variant reduced substantially, further confirming the role BRD4-BD2 in recruiting ILF3 to the locus (Fig. 9F).

Finally, we examined the effect of co-localization on Survivin at the transcriptional level. We isolated total RNA from HEK293T cells expressing NF110 in combination with wild type BRD4 or the non-functional variants N140A/ N433A. It is evident from qRT-PCR data that NF110 and BRD4 individually increased the expression of Survivin, albeit moderately, compared to non-transfected cells, and the NF110-BRD4 co-expression further enhanced the level of Survivin transcript, thus confirming their synergistic role in the locus (Fig. 9G). In contrast, BRD4 mutants reduced Survivin expression with N433A having a higher inhibitory effect, corroborating well with the BD2-mediated interaction between ILF3 and BRD4 to effect promoter localization and transcriptional activity. Furthermore, replacing wild type ILF3 with the acetylation-defective mutant prevented Survivin upregulation and cannot be rescued by wild type BRD4. Collectively, our results uncovered a novel mechanism, involving posttranslational acetylation on a highly conserved lysine residue in eukaryotic ILF3 and its recognition by BET bromodomain, to mediate expression of genes essential to cell division (Fig. 9H, I). The interaction between the two transcriptional regulators constitutes a mechanism for their localization to mitotic chromosomes and for regulation of cell cycle genes at a time when most transcription factors are displaced from chromatin. It is also interesting to note that the DZF domain of ILF3 is typically engaged with ILF2 (NF45) and the association is critical to posttranscriptional regulation of several mRNAs including Survivin.^28^ It is thus tempting to speculate that K100 acetylation in the DZF domain acts as an ‘interaction switch’ from ILF2 to BRD4. It would be of future interest to identify the deacetylases for K100ac mark and how the dynamic interplay between the writers (GCN5 and CBP), readers (BET), and erasers of ILF3 acetylation elicits signaling cascades in response to intrinsic and extrinsic stimuli.

## Conclusions

Specific interactions between acetyllysine and bromodomains, both of which are abundantly present in the human proteome, dictate critical cellular and developmental processes. In this study, we have tailored a chemoproteomic platform called IBPP to the closely related members of BET bromodomains BRD2-4 and BRDT, and characterized their distinct and shared interacting partners. Employing amber suppression mutagenesis, we introduced a photo-responsive unnatural amino acid into a highly conserved site within the aromatic cage of BRD2, 3 and BRDT without altering their biochemical integrity, and demonstrated the ability of the BET analogues to crosslink acetylated H4 upon photo-irradiation. Subsequent crosslinking with the acetylated proteome present in HEK293T cells along with affinity enrichment and systematic proteomic analysis revealed a set of human proteins as putative interacting partners of the bromodomains. The interactomes are implicated in essential cellular processes ranging from DNA replication and repair, transcription and RNA processing, protein folding and stability, and metabolic pathways, suggesting a multitude of potential functions for bromodomains beyond gene regulation. The interactome profiling revealed both shared and distinct interacting partners across the BET bromodomain. Our observations are consistent with the increasing number of reported interacting partners of BET bromodomains. Detailed *in vitro* binding study with synthesized protein segments carrying site-specific acetyllysine revealed that the bromodomains are indeed capable of recognizing proteins with canonical H4 motifs as well as distant sequence patterns. Analysis of eight high-resolution crystal structures of interactome-bound BRD3 and 4 complexes presented here highlights the unprecedented mode of association between the readers and the binding partners. Particularly notable was substitution of the second acetyllysine unit with a hydrophobic amino acid in peptide sequence that could fit into the conformationally flexible aromatic cage, thus establishing a critical recognition element in the expanded set of non-histone proteome for bromodomains. Subsequent biochemical investigations demonstrate how acetylation of a conserved lysine in a transcriptional regulator (ILF3) and its interaction with BET bromodomain (BRD4) regulates genes such as Survivin, a protein critical to mitotic cell division and an important anticancer drug target. Collectively, our biochemical, proteomic and structural findings strengthen the hypothesis that non-canonical sequence templates, beyond those residing in histones, regulate the majority of BET functions. We anticipate future studies to further validate the newly identified interactions in human cells and examine their potential mechanistic role in initiating downstream acetyllysine-mediated signaling cascades parallel to those regulated by protein phosphorylation and methylation. The structural studies presented here may serve as a platform for developing specific small-molecule inhibitors against distinct members of the interactome. Furthermore, the current findings will serve as a foundation for engineering orphan bromodomains whose interacting partners and physiological function have remained elusive.

### Methods

Synthetic methods and characterization of 4-azido-L-phenylalanine (AzF), additional methods, supplementary figures, NMR spectra, supplementary tables and associated references are given in Supplementary Information. Supplementary information is available in the online version of the paper.

## Acknowledgements

We thank the University of Pittsburgh, the Melanoma SPORE Program of the University of Pittsburgh Cancer Institute, the National Science Foundation (MCB-1817692), the National Institutes of Health (R01GM123234, R01GM130752 and P01GM118303) for financial support; the UPCI Cancer Biomarkers Facility (supported in part by award P30CA047904) for LC-MS/MS analysis; and Dr. D. Chakraborty and members of our laboratories for editing of the manuscript. We also thank the shared resource facilities at the Albert Einstein College of Medicine are supported in part by Cancer Center Grant P30 CA013330.

