## Supplementary methods, Figures and Tables for "Uncovering the Bromodomain Interactome using Site-Specific Azide-Acetyllysine Photochemistry, Proteomic Profiling and Structural Characterization": BET Supporting Information_bioRXIV.pdf

[†]Lead Author

**Table of Contents**

1. General materials, methods, and equipment S2
2. Plasmids, mutagenic primers, cell lines, and antibodies S2
3. Peptide synthesis and purification S3
4. Expression and purification of BET Bromodomains S4
5. Isothermal titration calorimetry S6
6. Photo-crosslinking with TAMRA peptide S6
7. Full length histone crosslinking and Western blotting S6
8. Cell Culture, transfection, cell lysis, and nuclear extraction S7
9. Immunoprecipitation S8
10. Gel Electrophoresis and Western Blotting of IP and Survivin lysates S9
11. RNA Extraction and qR-TPCR S10
12. Chromatin immunoprecipitation and quantitative PCR (ChIP-qPCR) S10
13. Photo-crosslinking with HEK293T cell lysate S11
14. LC-MS analysis of Expressed and Purified BET Proteins S11
15. LC-MS analysis of BET proteomics S12
16. Protein crystallization, data collection and analysis S13
17. Supplementary figures and tables S15
18. References S33

### 1. General materials, methods, and equipment

**Chemicals:** All chemicals were purchased from established vendors (e.g., Sigma-Aldrich, Acros Organics) and used without purification unless otherwise noted. Optima grade acetonitrile was obtained from Fisher Scientific and degassed under vacuum prior to use during HPLC purification. Analytical thin layer chromatography (TLC) was performed using EMD 250-micron flexible aluminum backed, UV $F_{254}$ pre-coated silica gel plates and visualized under UV light (254 nm) or by staining with phosphomolybdic acid, ninhydrin or anisaldehyde. Reaction solvents were removed by a Büchi rotary evaporator equipped with a dry ice-acetone condenser. Analytic and preparative HPLC was carried out on an Agilent 1220 Infinity HPLC with diode array detector. Concentration and lyophilization of aqueous samples were performed using Savant Sc210A SpeedVac Concentrator (Thermo), followed by Labconco Freeze-Dryer system.
Proton nuclear magnetic resonance spectra ($^1$H NMR) were recorded on Bruker Ultrashield$^{TM}$ Plus 600/500/400/300 MHz instruments at 24$^o$C. Chemical shifts of $^1$H and $^{13}$C NMR spectra are reported as δ in units of parts per million (ppm) relative to tetramethylsilane (δ 0.0) or residual solvent signals: chloroform-d (δ 7.26, singlet), methanol-$d_4$ (δ 3.30, quintet), and deuterium oxide-$d_2$ (δ 4.80, singlet). Coupling constants are expressed in Hz. Mass spectra were collected at the UPITT MASSSPEC lab on a Q-Exactive$^{TM}$ Thermo Scientific LC-MS with electron spray ionization (ESI) probe. Synthesis of p-Azidophenyl alanine (AzF) was accomplished following the reported method.[1]

### 2. Plasmids, mutagenic primers, cell lines and antibodies

All the plasmids for bacterial expression are obtained as gifts from individual laboratories or purchased from Addgene. Details of these constructs are given in Table S8. Mutagenic primers are obtained from Integrated DNA Technologies (Table S9). Competent bacterial cells used for protein expression and mutagenesis are mentioned in section 4. Human embryonic kidney 293T (HEK293T) cells are obtained from the American Type Culture Collection (ATCC) and used in the current study following manufacturer's protocol. All the antibodies used in the current study are purchased from established vendors and used following manufacturer's protocol.

### 3. Peptide synthesis and purification

Peptides **2-13** were synthesized by the University of Pittsburgh Peptide Synthesis Facility; Crude peptides were purified using preparative reversed-phase HPLC (XBridge C18, 5 µm, 10 x 250 mm column) eluting with a flow rate of 5 mL/min and a gradient of acetonitrile starting from 0% v/v to 50% v/v in 12 min and then to 100% v/v by 18 min in aqueous trifluoroacetic acid (0.1% v/v). Purified peptides were concentrated by SpeedVac concentrator then lyophilized. Dried peptides were resuspended in water and stored at -80 °C before use. Concentrations of peptides with no aromatic amino acids were determined based on the observation that 1 mg/ml peptide generates an absorbance value ($A_{205}$) of 30 at 205 nm. Concentrations of peptides with aromatic amino acids were determined based on the absorbance and extinction coefficient for the aromatic amino acid at lambda max, using molar ratio of amino acid to peptide to extrapolate. The integrity of the purified peptides was confirmed by MALDI mass spectrometry.

Peptides **14-15** were synthesized on solid support NovaPEG Rink Amide Resin (Sigma Aldrich #8550470005) in 5 mL fritted syringes. Amino acids were purchased from Sigma and used without further purification (Fmoc-Ala-OH #852003, Fmoc-Arg(Pbf)-OH #852067, Fmoc-Gly-OH #852001, Fmoc-Leu-OH #852011, Fmoc-Lys(Boc)-OH #852012, Fmoc-Lys(Ac)-OH #852042, Fmoc-Phe-OH #852016, Fmoc-Thr(tBu)-OH #852000). Resin was swelled 20 minutes with N,N-Dimethylformamide (DMF) (Acros #348430025) prior to addition of C-terminal amino acid. Peptides were prepared at a 0.05 mmol scale. Four equivalents of amino acid, four equivalents of O-(1H-6-Chlorobenzotriazole-1-yl)-1,1,3,3-tetramethyluronium hexafluorophosphate (HCTU) (Sigma #8.51012), and six equivalents of N,N-Diisopropylethylamine (Sigma #496219) were first dissolved in 1-Methyl-2-pyrrolidinone (Sigma #443778) via sonication with an Ultrasonic Cleaner water bath(VWR #97043-968). The amino acid mixture was added to the resin and stirred under atmospheric conditions at 250rpm for 45-60 minutes. Resin was then washed with DMF three times prior to addition of 20% 4-methylpiperidine in DMF (Sigma #M73206). Deprotection solution was mixed with resin for 10 minutes at 250 rpm two times. Resin was then washed with DMF three times, and the next amino acid was added. Once the N-terminal amino acid was deprotected, the resin was washed three times each with DMF, dichloromethane (DCM) (Acros #348465000), and methanol (Acros #364395000). Cleavage cocktail containing 91.5% TFA (Alfa Aesar L063374), 2.5% thioanisole (Acros T28002), 2.5% phenol (Alfa Aesar #J64011), 2.5%

deionized water, and 1% triisopropylsilane (Acros Organics # 214520100) was added to the resin. The sealed fritted syringe was rocked for 2 hours at room temperature using a fixed angle rocker (Fisher 07202202). After cleavage, the solvent was ejected from the syringe into a 50mL Falcon tube, leaving behind the resin in the syringe. The solution was condensed to remove roughly 50% TFA prior to precipitation via -80°C diethyl ether (Acros #326860010). Precipitate was then pelleted via centrifugation (Sorvall Legend XTR) at 3000 rpm for 5 mins. Supernatant was decanted. Crude peptide was the resuspended in 0.1% TFA prior to purification and characterization as described for peptides from the peptide facility. The MALDI-MS are provided in Supplementary Figures S2, 6 and Table S2.

### 4. Expression and purification of BET Bromodomain

The N-terminal 6xHis-tagged human BRD2 Bromodomain 1 bacterial expression construct pET was obtained from Vector Builder (cyagen). The N-terminal 6xHis-tagged human BRD3, BRD4 and BRDT Bromodomain 1 bacterial expression constructs pNIC28-Bsa4 were obtained from addgene (#38940, #38942, and #38898, respectively). Wild type plasmids were transformed into *E. coli* BL21 (DE3) Star competent cells (Invitrogen) using pNIC28-Bsa4 kanamycin-resistant vector.[1] A single colony was added to 10 mL Luria-Bertani (LB) Miller broth with 50 µg/mL kanamycin sulfate (BRD3, BRD4, BRDT) or 100 µg/mL ampicillin (BRD2), and grown overnight at 37 °C. Overnight cultures were diluted 100-fold the following morning and allowed to grow at 37 °C to an optical density (OD600) of 0.8, at which time 0.35 mM IPTG was added to induce protein expression overnight at 17 °C with in an Innova 44® Incubator shaker (New Brunswick Scientific). Protein purification was performed as follows: Cells were harvested by centrifugation for 20 minutes at 4000 rpm, and supernatant was discarded. Resulting cell pellets were resuspended in 15 mL lysis buffer (50 mM Tris-HCl pH 8.0, 200 mM NaCl, 5 mM β-mercaptoethanol, 10% v/v glycerol, 25 mM imidazole, 1 mg/mL Lysozyme, 10µg/mL or 20U/mL DNase, and Roche protease inhibitor cocktail), then lysed by pulsed sonication (Qsonica-Q700, Fisherbrand Replaceable Microtip 1/8" Probe, pulsed 10 seconds on, 10 seconds off, for 2 minutes processing time, 60Amps; repeated sonication a second time if solution was still opaque and viscous) at 13000 rpm, 19632.4 x g for 40 min at 4 °C. Soluble extracts were subjected to Ni-NTA agarose resin (Thermo) according to manufacturer's instructions. Bound protein was washed by allowing supernatant to flow through the column followed by 20 volumes of washing buffer (50 mM Tris-HCl pH 8.0, 200 mM NaCl, 5 mM β-mercaptoethanol, 10% v/v glycerol, and 25 mM imidazole). Proteins were then eluted with elution buffer (50 mM Tris-HCl pH 8.0, 200 mM NaCl, 5 mM β-mercaptoethanol, 10% v/v glycerol, and 400 mM imidazole). Eluted proteins were then further purified by size exclusion chromatography (Superdex-75) using AKTA pure FPLC system (GE healthcare) with FPLC buffer (50 mM HEPES pH 7.5, 150 mM NaCl, and 0-5% v/v glycerol). Purified protein was concentrated using Amicon Ultra-10k centrifugal filter device (Merck Millipore Ltd.), at which time protein concentration was determined using Bradford assay kit (BioRad Laboratories) with BSA as a standard. Concentrated proteins were stored at -80°C before use. BET TAG variants were generated using QuikChange Lightning site-directed mutagenesis kit (Agilent Technologies). The resulting mutant plasmids were confirmed by DNA sequencing.

To express BRD2-L108AzF, BRD3-L68AzF, BRD4-L92AzF, and BRDT-L61AzF, BL21 Star (DE3) cells were co-transformed with pEVOL-based *M. jannaschii* TyrRS-$tRNA_{CUA}^{Tyr}$ pair for AzF (addgene ID: 31186).[2-3] Cells were recovered for 1 hour and 45 minutes in 200 µL SOC medium in a 37°C shaker prior to plating on an LB Miller agar plate containing 50 µg/mL kanamycin sulfate or ampicillin and 35 µg/mL chloramphenicol. Single colonies were picked and added to each of four inoculates containing 10 mL of Luria-Bertani (LB) Miller broth in presence of appropriate antibiotics (35 µg/mL chloramphenicol and 50 µg/mL kanamycin sulfate (BRD3, BRD4, BRDT) or 100 µg/mL ampicillin (BRD2)). Overnight cultures were centrifuged (ThermoScientific Sorvall Legend XTR Centrifuge, TX-1000 rotor, 4°C) for 10 min at 1000 x g, 2100 rpm. 9 mL of LB Miller broth was removed and the cell pellet was resuspended in the remaining 1mL LB Miller broth and used to inoculate 1L of GMML medium (M9 minimal media supplemented with 1% v/v glycerol, 300 µM leucine, 1 mM $MgSO_4$, 0.1 mM $CaCl_2$, 50 µg/mL kanamycin sulfate, and 35 µg/mL chloramphenicol, and trace amounts of $Na_2MoO_4$, $CoCl_2$, $CuSO_4$, $MnSO_4$, $MgSO_4$, $FeCl_2$, $CaCl_2$, and $H_3BO_3$). Cells were allowed to grow at 37 °C to an optical density (OD600) of 0.8. AzF **1** was prepared by diluting in 20mL sterilized deionized water and added aseptically to a final concentration of 1mM. Cells were allowed to shake an additional 30 minutes at 17 °C, at which time the synthetase expression was induced with 0.05% w/v arabinose and allowed to shake an additional 30 minutes at 17 °C. Finally, 0.425 mM IPTG was added to induce BET protein expression while shaking overnight at 17 °C with an Innova 44® Incubator shaker (New Brunswick Scientific). Protein purification, concentration, and storage were performed as previously described.

### 5. Isothermal titration calorimetry

Isothermal titration calorimetry (ITC) was performed with an $ITC_{200}$ instrument (MicroCal, Malvern). Experiments were conducted at 15 °C while stirring at 750 rpm. Buffers of protein and peptides were matched to 50mM HEPES pH 7.5 and 150 mM NaCl. Each titration was performed as follows: one initial injection of 0.4 µL for 0.8 seconds, followed by 19 injections of 2.0 µL for 4 seconds, with ≥2 mins between each injection. The initial injection was discarded prior to data analysis. The microsyringe (40µL) was loaded with 1-6mM peptide and injected into the cell (200µL), occupied by a BET protein at a concentration of 100 to 200 µM. Data was fitted to a single binding site model using Microcal $ITC_{200}$ Software with Origin Lab 7 to yield enthalpies of binding (ΔH) and binding constants ($K_a$). Further thermodynamic parameters (changes in entropy ΔS, changes in free energy ΔG, and dissociation constants ($K_d$)) were calculated from these values. The thermodynamic parameters are provided in Table S3.

### 6. Photo-crosslinking experiment with peptides and in-gel fluorescence

For peptide photo-crosslinking experiments, 1 µM TAMRA-labeled tetra-acetylated H4 peptide **3** was preincubated with 25 µM of wild type BRD2/3 or variant in a buffer containing 10 mM Tris-HCl pH 7.5, 150 mM NaCl, 0.05% Tween 20, and 0.5 mM TCEP for 30 minutes at room temperature in darkness. Samples were then kept 3 inches underneath a UV lamp (MAXIMA ML-3500S UV, 50,000 µW/cm$^2$, 350nm-385nm 50% irradiated, 365nm 100% irradiated) for 30 minutes at 4 $^{o}$C on ice. Negative controls were kept in a drawer on ice for the full 30 minutes. 10uL of sample was mixed with 10uL 4X Laemmli dye (Biorad #1610747) and separated on a 4-12 % Criterion XT precast gel (Bio-Rad Laboratories). Gels were imaged on a ChemiDoc MP Imaging system using TAMRA fluorophore excitation wavelength (Emission filter 605/50, Light: green Epi illumination). The gel was subsequently stained with Coomassie brilliant blue R-250 staining solution to confirm the presence of proteins in all the samples and again imaged with the ChemiDoc. For inhibition studies, BRD2/3 variants were preincubated with 5 µM of JQ1 for 30 min prior to the preceding protocol.

### 7. Full length histone crosslinking and Western blotting

Photo-crosslinking experiments were conducted as follows: 10µM acetylated histone (H4$K_C$5Ac/ H4$K_C$8Ac/ H4$K_C$12Ac/ H4$K_C$16Ac)[4] or ~0.5ug histone H4 (prepared as histone extracts as described in reference 5 and using BSA standards to estimate concentration)[5] and 50μM BRD2 L108AzF or BRD3 L68AzF were diluted to 20μL with buffer (10mM Tris HCl pH 7.5, 150mM NaCl, 0.05% w/v TWEEN 20, and 0.5mM TCEP). Vortexed (Fisher Scientific Mini Vortexer 120V) and centrifuged (Fisherbrand Sprout) samples were split into two 10μL aliquots. Samples were preincubated and crosslinked as described above for two timepoints of 0 and 30-60 minutes. Samples were diluted 1:1 with 10μL of 4X Laemmli loading buffer, incubated at 95°C for 10 minutes, and loaded onto a precast 4-12% w/v Bis-Tris polyacrylamide gel (BIORAD cat#3450123). The protein (crosslinked and free) bands were separated by electrophoresis at 150V for 1 hour.

Separated protein bands were then transferred to a 0.2 μm nitrocellulose membrane (BIORAD cat #1620112) at constant voltage of 80V for 1 hour. The membrane was washed once with TBST (50 mM Tris HCl pH 7.4, 150 mM NaCl, 0.01% w/v Tween-20) then blocked with 5% w/v milk in TBST buffer for H4 and His H-3 or 5% w/v BSA in TBST buffer for 6xHis for 1 hour. Immunoblotting was performed with 1:500 to 1:300 diluted H4 primary antibody (Histone-H4 mAb, cat# 61521, Active Motif) or 1:300 to 1:200 diluted His primary antibody (His-probe (H-3) mAb, cat# sc-8036, Santa Cruz Biotechnology) or 1:300 diluted His primary antibody (Anti-6X His tag, cat# ab9108, abcam) and incubated overnight at 4 °C. Antibody solutions were removed, then membranes were washed with TBST three times. Secondary antibody was added to membranes at a dilution of 1:5000 antimouse (HRP Goat anti-Mouse IgG, cat #15014, Active Motif) for H4 and His H-3 or 1:5000 antirabbit (Anti-rabbit IgG, HRP-linked Antibody, cat #7074S, Cell signaling technology) and incubated at room temperature for ≥1 hour. Membranes were then washed three times with TBST, and then visualized using VISIGLO HRP Chemiluminescent substrates A and B (cat# N252-120ML and N253-120ML, aMReSCO) following manufacturer's protocol.

### 8. Cell Culture, transfection, cell lysis, and nuclear extraction

Human embryonic kidney (HEK) 293T cells were grown in Dulbecco modified Eagle medium (DMEM) (Gibco) supplemented with 10% fetal calf serum, 1x Penicillin-Streptomycin-Amphotericin in a humidified atmosphere containing 5% $CO_2$ in appropriately sized flask or dish pretreated for adherent cell lines (6-well for qRT-PCR and western blotting, T75 for immunoprecipitation, 150mm dish for ChIP). Cells were transfected at 60-80% confluency with a ratio of 1 microgram DNA per 1.2 x $10^6$ cells for BRD4, hnRNPK, and ILF3 or 1.25 micrograms DNA per 1.2 x $10^6$ cells for CBP and GCN5. A 3:1 ratio of Lipofectamine 2000 (Invitrogen) to DNA was used. Both DNA and lipofectamine were incubated separately for five minutes at room temperature in OptiMEM (Gibco) before combining and incubating at room temperature for a minimum 20 minutes. Six hours after transfection, media was replaced and (if applicable, eg IP studies) suberoylanilide hydroxamic acid (SAHA) (Cayman Chemicals Company) was added to a final concentration of 3 uM. Cells were collected a minimum 24 hours post transfection.

Cell lysis: To lyse HEK cells, cell pellets were resuspended in RIPA buffer (Alfa Aesar #J61529) supplemented with 5 mM TCEP, 1X Roche protease inhibitor cocktail, and 1 mM PMSF. Resuspended cells were incubated on ice for ten minutes prior to sonication (Qsonica-Q700, Cuphorn, pulsed 1 minute on, 30 seconds off, for 5 minutes processing time, 100 Amps). Insoluble cell debris were pelleted via centrifugation at 16000rpm for 10 minutes at 4°C (Eppendorf® Microcentrifuges, 5424R). Supernatant was transferred to a new microcentrifuge tube for Bradford quantification and gel electrophoresis.

Nuclear Extraction: To lyse HEK cells, cell pellets were resuspended in nuclear isolation buffer (60mM potassium chloride, 15mM sodium chloride, 5mM magnesium chloride, 1mM calcium chloride, 1 mM Dithiothreitol, 2 mM sodium vanadate, 250 mM sucrose, 1X Roche protease inhibitor cocktail, 0.3% NP40, 1mM PMSF, and 15mM Tris pH 7.5) and incubated on ice 5 minutes. Nuclei were then pelleted via centrifugation at 2000 rcf for 5 minutes at 4°C (Eppendorf® Microcentrifuges, 5424R). Nuclear isolation buffer was removed via aspiration. Nuclei were then resuspended in nuclear extraction buffer (150 mM sodium chloride, 1 mM EDTA, 5% Glycerol, 1X Roche protease inhibitor cocktail, 0.2% NP40, 60 mM PMSF, and 25mM Tris pH 8). Resuspended nuclei were lysed via sonication (Qsonica-Q700, Cuphorn, pulsed 1 minute on, 30 seconds off, for 5 minutes processing time, 100 Amps). Nuclear debris were pelleted at 2000 rcf for 5 minutes at 4°C, and supernatant was transferred to a new microcentrifuge tube for Bradford quantification and immunoprecipitation or gel electrophoresis.

### 9. Immunoprecipitation

Immunoprecipitation was performed with a Dynabeads Protein A Immunoprecipitation kit (Invitrogen 10006D) according to the manufacturer protocol in publication MAN0017347, with the following adjustments: 1. Dynabeads were incubated with antibody (1:50 uL of BRD4 (E2A7X) Rabbit mAb CST 13440S, 1:20 uL of NF90/NF110 Rabbit anti-Human, Polyclonal, Bethyl Laboratories A303-120A, 1:50 hnRNP K (D9A8) Rabbit mAb CST 9081S, 1:50 NF90 mouse Santa Cruz biotech sc-377406, or same micrograms antibody IgG (Rabbit (DA1E) mAb IgG XP Isotype Control CST 3900S as micrograms of test antibody) for 2 hours at 4°C with rotation in Ab Binding & Washing solution. 2. No crosslinking of antibody was performed. 3. After pelleting and washing beads, nuclear extract samples were added to beads in a total of 200 uL nuclear extraction buffer with an equal number of micrograms per sample (200-1000ug). 4. Beads and nuclear extracts were incubated overnight at 4°C with rotation. 5. Only two washes were performed with Washing Buffer. 6. Samples were eluted with 20uL of Elution Buffer and 10 uL of 4X Laemmli Dye (Bio-Rad).

**10. Gel Electrophoresis and Western Blotting of IP and Survivin lysates**

Samples were prepared with 4X Laemmli Dye (Bio-Rad) and loaded onto 4–12% Criterion™ XT Bis-Tris protein gels (Bio-Rad #3450123) and subjected to electrophoresis (Criterion Precast Tank, Bio-Rad 1656001) at 150V for 1 hour in 1X MES buffer (prepared from Invitrogen™ Novex™ 20X Bolt™ MES SDS Running Buffer). After electrophoresis, gels were removed from precast cassettes and incubated for 5 minutes with semidry transfer buffer (48mM Tris Base, 39 mM glycine, 0.0375% SDS, 20% methanol) along with two extra-thick blotting papers and 0.45 µm nitrocellulose membrane (BIORAD cat #1620112) for IP or 0.2 µm PVDF (Immobilon-$P^{SQ}$ ISEQ00010) membrane for Survivin blots preactivated with methanol for 30 seconds. Semidry blotting was performed at a constant amplitude of 5.5 mA per square centimeter for 30 minutes with a maximum voltage of 25 V (Bio-Rad #1703940). After transfer, the membrane was washed once with TBST (50 mM Tris HCl pH 7.4, 150 mM NaCl, 0.01% Tween-20) prior to blocking with 5% milk or 5% BSA (according to vendor's recommendation for primary antibody) in TBST buffer for 1 hour.

Immunoblotting was performed with 1:1000 primary antibody for BRD4, hnRNPK, Flag, Survivin, or β Actin or 1:2000 NF90/110 or 1:500 anti-acetyllysine (Acetylated Lysine Rabbit anti-All, Invitrogen PIMA533031) and incubated overnight at 4 °C. Antibody solutions were then removed, and membranes were washed with TBST three times. Secondary antibody was then added to membranes, either 1:5000 anti-rabbit (Goat Anti-Rabbit IgG-HRP, cat #4030-05,

Southern Biotech) for Survivin or 1:5000 anti-mouse (HRP Goat anti-Mouse IgG, cat #15014, Active Motif) for β actin, or 1:1000 Protein-A-HRP (12291S CST) for immunoprecipitation blots and incubated at room temperature for 1 hour. Secondary antibody was washed off three times with TBST, and membranes were visualized using chemiluminescence (Thermo Scientific™ Pierce™ ECL Western Blotting Substrate PI32106) following manufacturer's protocol.

### 11. RNA Extraction and qRT-PCR

Total RNA was extracted from HEK cell pellets as described by manufacturer protocol (E.Z.N.A.® HP Total RNA Kit, Omega Bio-tek® - R6812-02, HP Total RNA Kit, VWR, 101414-852). RNA concentration was measured via the RNA program on NanoDrop OneC. One microgram of total RNA from each sample was used to generate cDNA via manufacturer protocol (cDNA Supermix qScript, VWR #101414-106). The resulting PCR product was diluted 1:1 with nuclease free water. 96-well plates (ThermoScientific #AB2800) were prepared with 2 µL 5 µM forward primer and 5 µM reverse primer in appropriate wells. Master mixes were prepared for triplicate qRT-PCR containing 1 µL cDNA, 5 µL nuclease free water, and 10 µL Sybr Green (PerfeCTa® SYBR® Green SuperMix Reaction Mixes, QuantaBio, VWR# 101414-152). 16 µL of the master mix was added to appropriate wells. Plates were then microsealed (Microseal® 'B' PCR Plate Sealing Film, adhesive, optical BioRad #MSB1001) and centrifuged at 3000 rpm for 3 minutes at 4°C. qPCR was performed using a CFX96 Real-Time PCR Detection System (Bio-Rad) with an initial denaturation step of 95°C followed by 55 cycles of 10 seconds at 95°C then 30 seconds at 55°C. Qt values of less than 30 were considered valid.

### 12. Chromatin immunoprecipitation and quantitative PCR (ChIP-qPCR)

Chromatin digestion, analysis, and quantification was performed for transfected 15 cm dishes as described by manufacturer protocol [SimpleChIP® Plus Enzymatic Chromatin IP Kit (Magnetic Beads) CST #9005]. Chromatin immunoprecipitations with 15 µL of NF90/NF110 antibody and 10 µg chromatin and subsequent DNA purification were performed as per manufacturer protocol. qPCR was performed as described for cDNA in Section 11, with the adjustment of 2 µL of ChIP DNA added and 4 µL nuclease free water in the master mixes. The primers for qRT-PCR and ChIP-qPCR are provided in Table S10.

**13. Photo-crosslinking with HEK293T cell lysate**

For photo-crosslinking studies, 2.5-3.0 mg of SAHA treated HEK293T cell lysates were incubated with 50 μM of the AzF variants of BRD2, 3 and BRDT in a buffer containing 10 mM Tris-HCl pH 7.5, 150 mM NaCl, 0.05% Tween 20, and 0.5 mM TCEP. After 1 h of incubation at room temperature, the samples were subjected to UV irradiation at 365 nm for 30 min at 4°C. Negative controls were not subjected to UV exposure. Samples were then bound to Ni-NTA agarose resin and incubated for 1 hr. at 4°C with gentle rotation. To remove un-crosslinked proteins, present in cell lysates, samples were washed with washing buffer (50 mM Tris-HCl pH 8.0, 400 mM KCl, 5% Triton X-100). During each washing step samples were incubated at 60°C for 5 min. Finally, the proteins were eluted with a buffer containing 50 mM Tris-HCl pH 8.0, 150 mM NaCl, 5 mM *β*-mercaptoethanol, and 400 mM imidazole. The eluted proteins were separated on a 4-12% Criterion XT precast SDS-PAGE gel (Bio-Rad Laboratories) and analyzed by Western blotting and tandem mass spectrometry.

**14. LC-MS analysis of Expressed and Purified BET Proteins**

For each HRMS, 5 μL of 5 μM protein was injected into a Thermo Scientific Q-Exactive Oribtrap. LCMS was run on a Thermo ProSwift RP-2H analytical 4.6x50mm SS column with ESI positive mode for 30 minutes with a gradient of 26% to 80% acetonitrile in 0.1% formic acid. Flow rate was set to 200 μL/minute. Chromatogram peak width was 10 seconds, resolution was 17,500, microscans was 5, maximum IT was 250 s, scan range was 500-3000 m/z, and AGC target was 3 x $10^6$. After LCMS, data was analyzed using Thermo Scientific Protein Deconvolution 3.0 software via Manual ReSpect (isotopically unresolved) method (chromatogram parameters: high sensitivity, auto spectral averaging False) (main parameters:, negative charge false, charge carrier $H^+$ (1.00727663), m/z range 1000-3000, output range 10000-25000, mass tolerance 5 ppm, charge state range 10-100, calculate XIC True, Peak Model Intact Protein) (advanced parameters: minimum peak significance 1 standard deviation, Noise Rejection 95% Confidence, Use relative Intensities True, number of iterations 3, minimum adjacent charges 6-10, number of peak modes 1, resolution at 400m/z 12374, and left/right peak shape 2:2).

**15. LC-MS/MS analysis of BET Proteomics.[6]**

*In gel trypsin digestion.* In gel trypsin digestion was carried out as previously described (1). Excised gel bands were washed with HPLC water and destained with 50% acetonitrile (ACN)/25mM ammonium bicarbonate until no visible staining. Gel pieces were dehydrated with 100% ACN, reduced with 10mM dithiothreitol (DTT) at 56°C for 1 hour, followed by alkylation with 55mM iodoacetamide (IAA) at room temperature for 45min in the dark. Gel pieces were then again dehydrated with 100% ACN to remove excess DTT and IAA, and rehydrated with 20ng/µl trypsin/25mM ammonium bicarbonate and digested overnight at 37°C. The resultant tryptic peptides were extracted with 70% ACN/5% formic acid, vacuum dried and re-constituted in18µl 0.1% formic acid.

*Tandem mass spectrometry.* Proteolytic peptides from in gel trypsin digestion were analyzed by a nanoflow reverse-phased liquid chromatography tandem mass spectrometry (LC-MS/MS). Tryptic peptides were loaded onto a C18 column (PicoChip™ column packed with 10.5cm Reprosil C18 3µm120Å chromatography media with a 75µm ID column and a 15µm tip, New Objective, Inc., Woburn, MA) using a Dionex HPLC system (Dionex Ultimate 3000, ThermoFisher Scientific, San Jose, CA) operated with a double-split system to provide an in-column nano-flow rate (~300nl/min). Mobile phases used were 0.1% formic acid for A and 0.1% formic acid in acetonitrile for B. Peptides were eluted off the column using a 52-minute gradient (2-40% B in 42 min, 40-95% B in 1min, 95% B for 1 min, 2% B for 8 min) and injected into a linear ion trap MS (LTQ-XL, ThermoFisher Scientific) through electrospray.

The LTQ XL was operated in a date-dependent MS/MS mode in which each full MS spectrum [acquired at 30000 automatic gain control (AGC) target, 50ms maximum ion accumulation time, precursor ion selection range of m/z 300 to1800] was followed by MS/MS scans of the 5 most abundant molecular ions determined from full MS scan (acquired based on the setting of 1000 signal threshold, 10000 AGC target, 100ms maximum accumulation time, 2.0 Da isolation width, 30ms activation time and 35% normalized collision energy). Dynamic exclusion was enabled to minimize redundant selection of peptides previously selected for CID.

*Peptide identification by database search.* MS/MS spectra were searched using MASCOT search engine (Version 2.4.0, Matrix Science Ltd) against the UniProt human proteome database. The following modifications were used: static modification of cysteine (carboxyamidomethylation, +57.05 Da), variable modification of methionine (oxidation, +15.99Da). The mass tolerance was set at 1.4Da for the precursor ions and 0.8 Da for the fragment ions. Peptide identifications were filtered using PeptideProphet™ and ProteinProphet® algorithms with a protein threshold cutoff of 99% and peptide threshold cutoff of 90% implemented in Scaffold™ (Proteome Software, Portland, Oregon, USA). The analyzed proteomic data are provided in Table S4-6.

**16. Protein crystallization, data collection and analysis**

***Protein crystallization and crystal harvest:*** Initial screening of domain 1 of BRD3 and BRD4 proteins in complex with acetylated peptides were performed using 800 nl (protein:mother liquor=1:1) sitting drops at a concentration between 12-15 mg/ml (~1-1.3 mM) with 2.5 molar excess of peptides with a Crystal Gryphon (Art Robbins Instruments, USA) and utilizing Index HT and Crystal Screen HT (Hampton Research). Crystallization conditions and ligand identities for individual complex are summarized in Table S7. For data collection at synchrotron, crystals were cryo-preserved by addition of either ethylene glycol or glycerol to the mother liquor prior to flash freezing in liquid nitrogen using appropriately sized micro-loop.

***Data collection:*** Cryo-preserved crystals were screened on ID-31 (LRL-CAT) beamline at the Argonne National Laboratory (APS) or on ID-17 (AMX/FMX) beamline at the Brookhaven National Laboratory (BNL) (Table S7). Complete data sets were collected on crystals, which exhibited good quality diffraction patterns. Data were collected on a CCD Rayonix MX-225 HE detector (APS ID-31) or Eiger 9M (AMX) and Eiger 16M (FMX) detectors.

***Data processing, structure determination, model building, refinement and analysis:*** Data from single crystals were collected using MOSFLM[7] (APS ID-31) or LSDCGuiREMOTE (BNL ID-17), integrated and scaled using either HKL-3000[8] and AIMLESS[9]. Initial phases of the domains were determined by molecular replacement using PHASER[10] with refined coordinates of the domains (BRD3: RCSB ID 2NXB and BRD4: RCSB ID 6DL2) as search models. After determining the phases an atomic model was built into the density using automated model building program BUCCANEER[11] and manually inspected using COOT[12]. The model was refined with REFMAC5[13] and PHENIX[14-15]. All crystals of BRD3 complexes contained two molecules of domain 1 in the asymmetric unit (protomers 1/chain A and 2/chains B). For structural interpretation of the complexes protomer 1 (chain A). No appreciable electron density corresponding to the acetylated ILF3 segment was observed for protomer 2 (chain B) of the BRD3-ILF3 complex. All crystals of BRD4 complexes contained one molecules of the domain in the asymmetric unit (chain A). Analyses of the structures were performed in COOT and MOLPROBITY[16]. Crystallographic statistics and RCSB accession codes are provided in Table S7.

### 17. Supplementary figures and tables

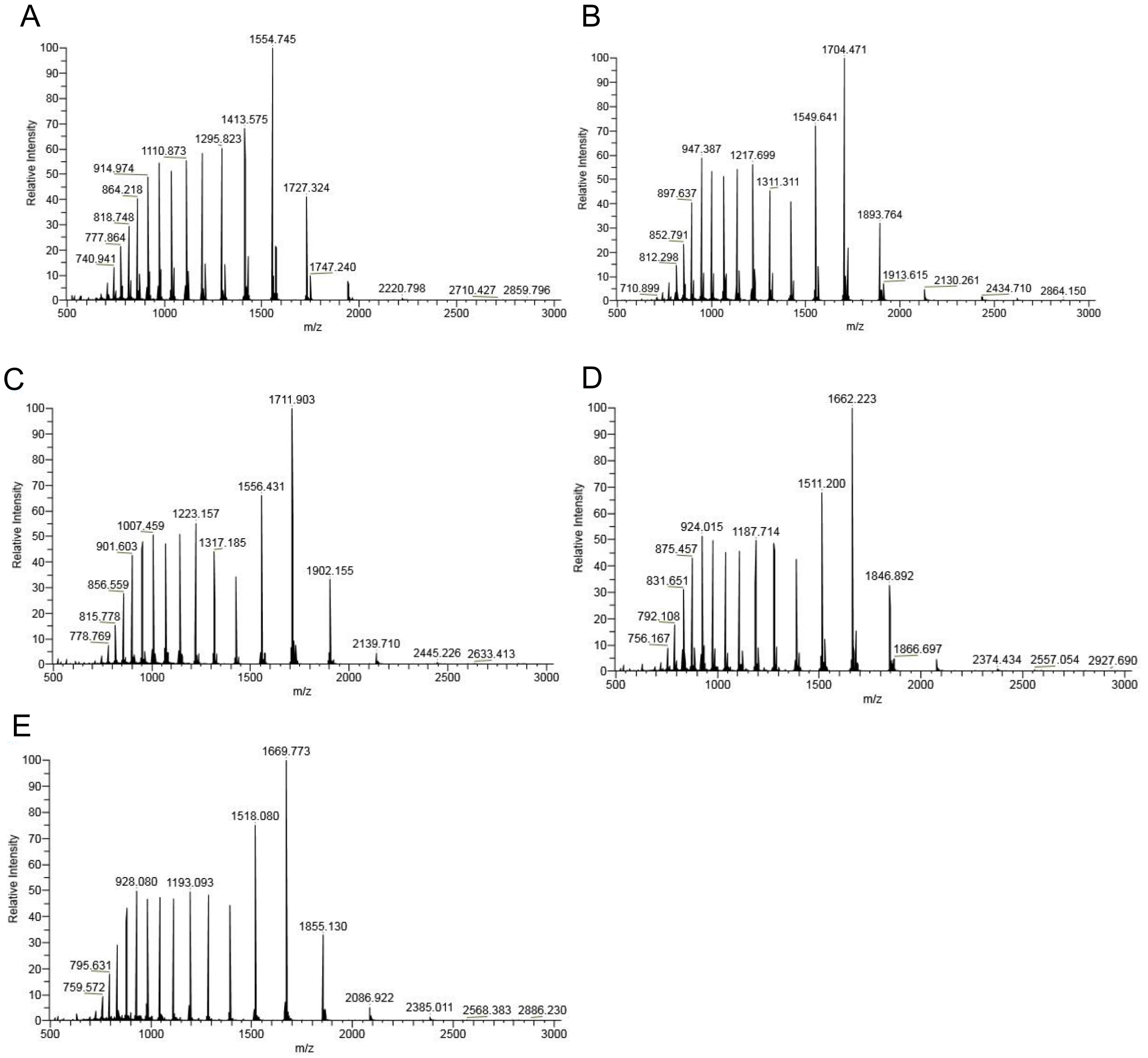


**Supplementary Figure S1:** HRMS Average Mass Protein Deconvolution Data: (A) WT BRD2-BD1; (B) WT BRD3-BD1; (C) BRD3-BD1 L68AzF; (D) WT BRDT-BD1 and (E) BRDT-BD1 L61AzF (E). See Supplementary Table S1 for expected and observed masses.

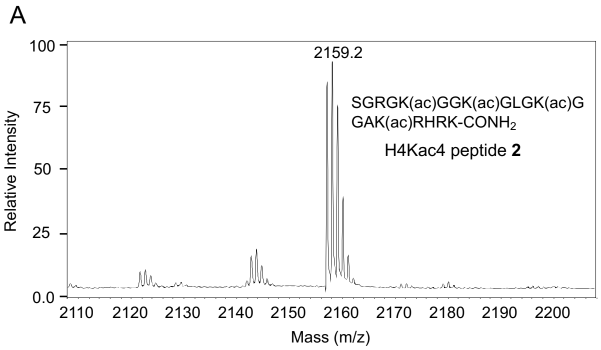


**Supplementary Figure S2:** MALDI-MS spectra of H4Kac4 peptide **2** (A) and TAMRA-H4Kac4 peptide **3** (B).

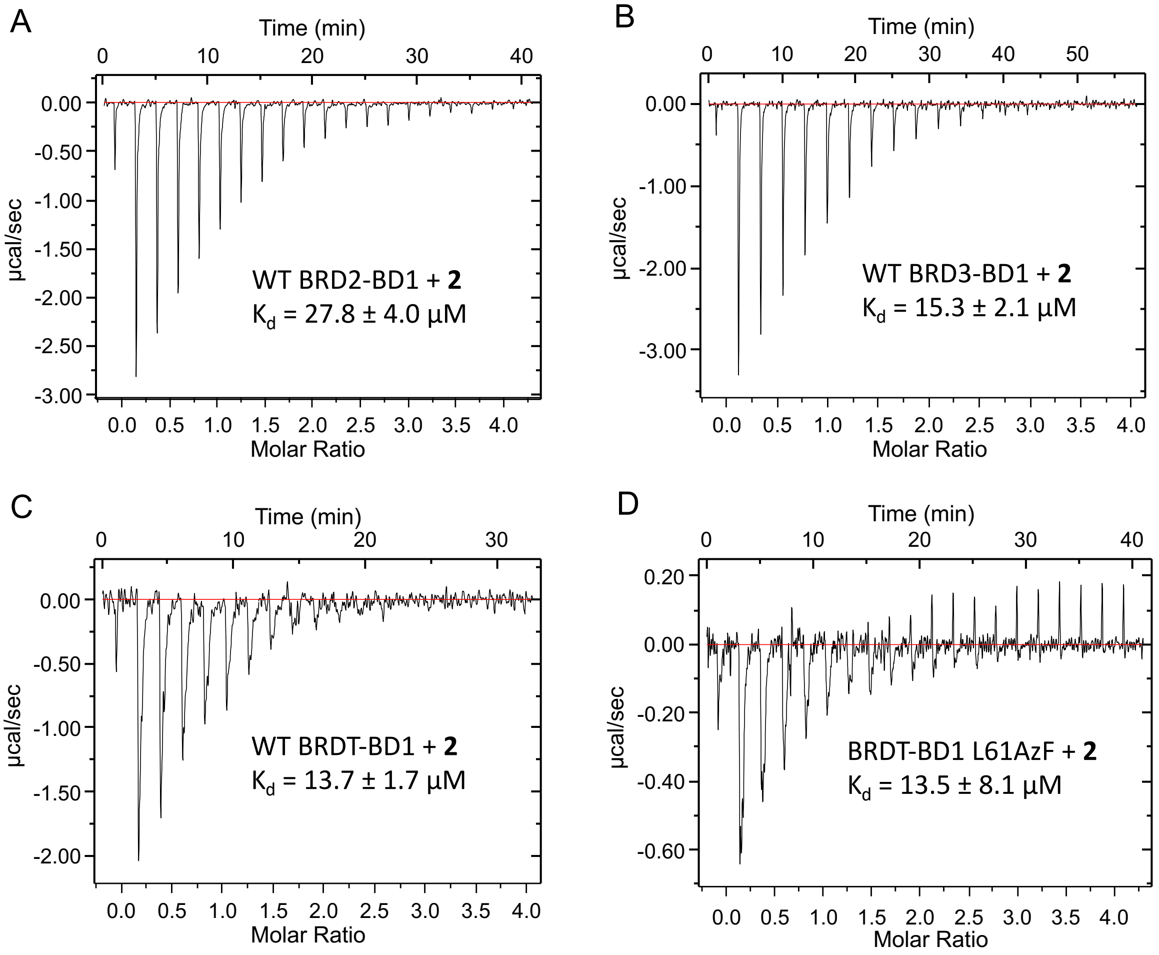


**Supplementary Figure S3:** Binding isotherms of BET proteins with H4Kac4 peptide **2**: (A) WT BRD2-BD1; (B) WT BRD3-BD1; (C) WT BRDT-BD1 and (D) BRDT-BD1 L61AzF as determined by isothermal titration calorimetry with H4Kac4 peptide **2.**

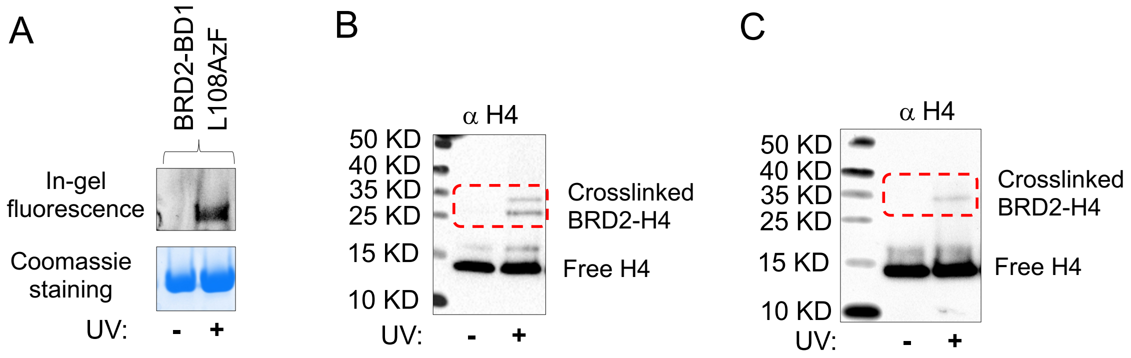


**Supplementary Figure S4:** Photo-crosslinking of BRD2-BD1 L108AzF with H4 peptide and full-length proteins. (A) Upon incubation with peptide **3**, samples were exposed to 365 nm UV light followed by in-gel fluorescence using 532 nm light ($\lambda_{max}$ for TAMRA). Samples that underwent successful photo-crosslinking upon exposure to 365 nm UV light are indicated by bands visible under 532 nm light as shown as in-gel fluorescence. Coomassie staining of the same gel showed presence of proteins in all the samples. (B) Visualization of the crosslinking of H4$K_C$5ac to BRD2-BD1 L108AzF using anti-H4 antibody. Successful crosslinking was observed when the sample was exposed to UV light. (C) Visualization of crosslinked endogenous H4 isolated from HEK293T cells to BRD2-BD1 L108AzF using anti-H4 antibody. Successful crosslinking was observed when the sample was exposed to UV light.

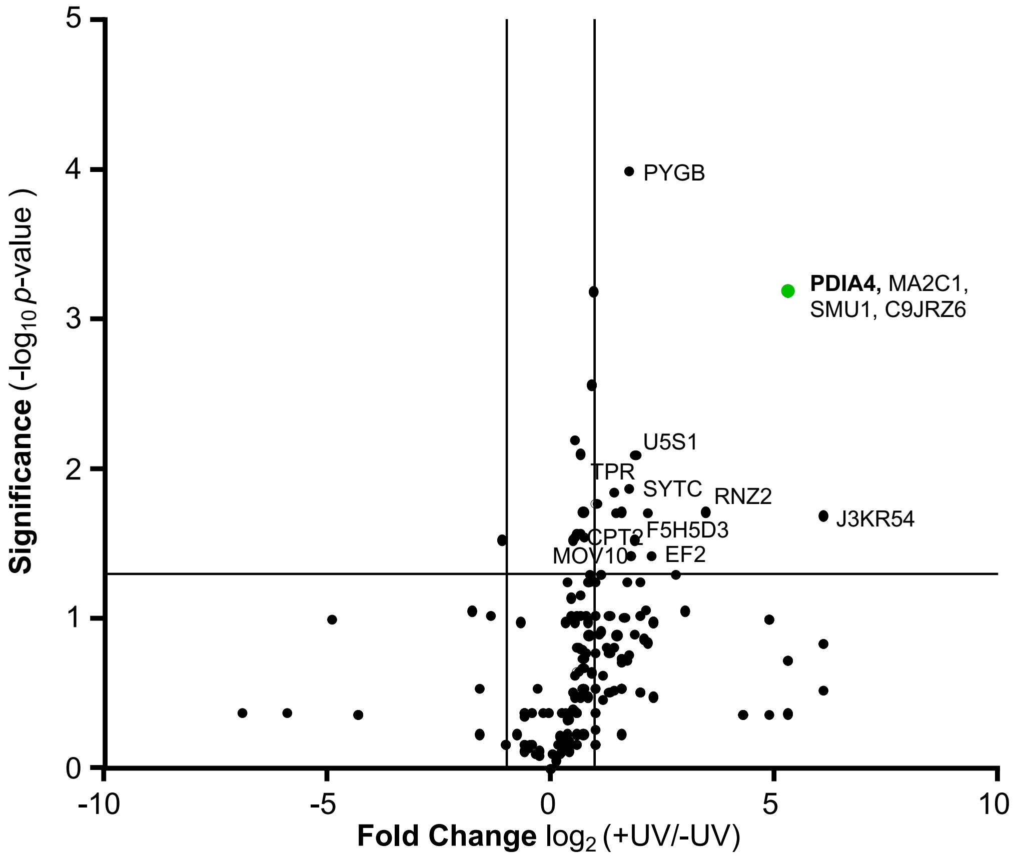


**Supplementary Figure S5:** Volcano Plot represents potential interacting partners of BRDT-BD1 identified using proteomic analysis.

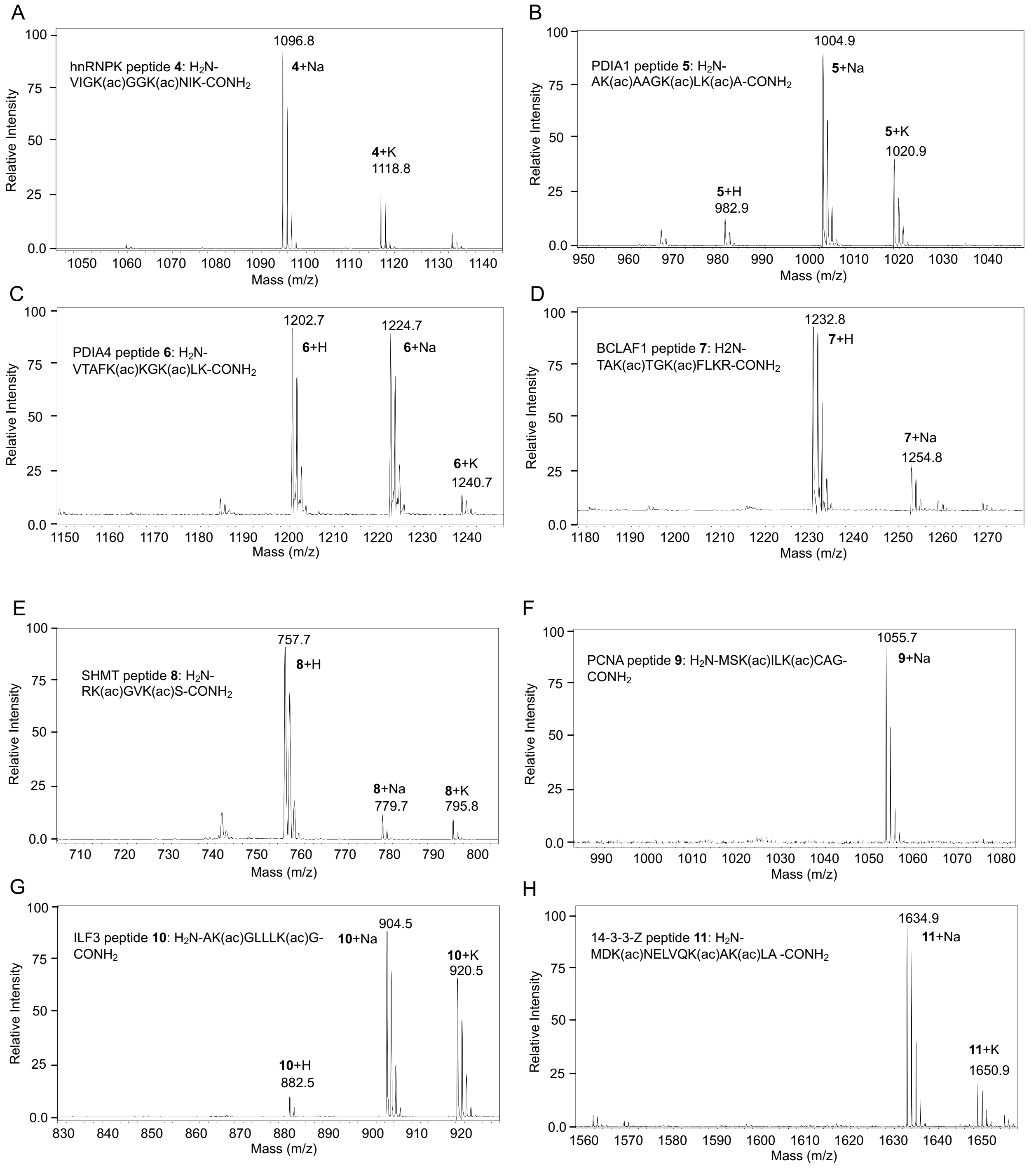


**Supplementary Figure S6:** MALDI-MS spectra of Peptides used for ITC: (A) hnRNPK **4**, (B) PDIA1 **5**, (C) PDIA4 **6**, (D) BCLAF1 **7**, (E) SHMT **8**, (F) PCNA **9**, (G) ILF3 **10**, (H) 14-3-3-Z **11**.

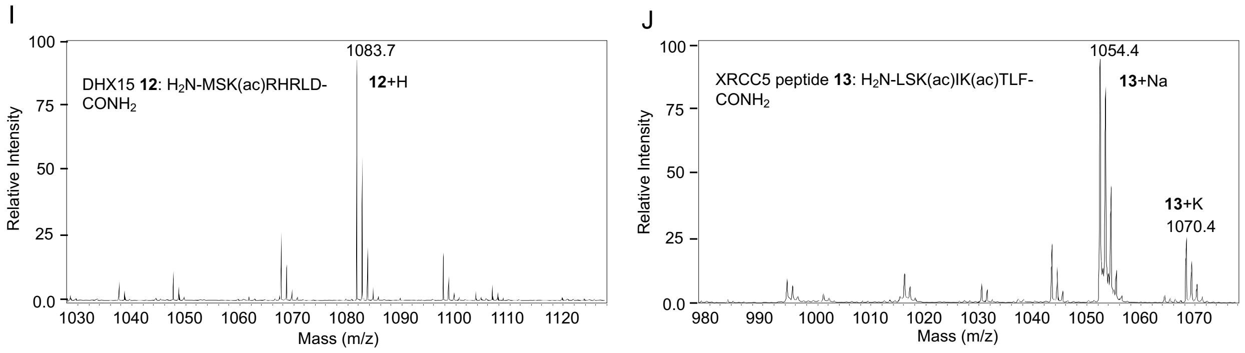


**Supplementary Figure S6 cont'd:** MALDI-MS spectra of Peptides used for ITC: (I) DHX15 **12**, (J) XRCC5 **13**.

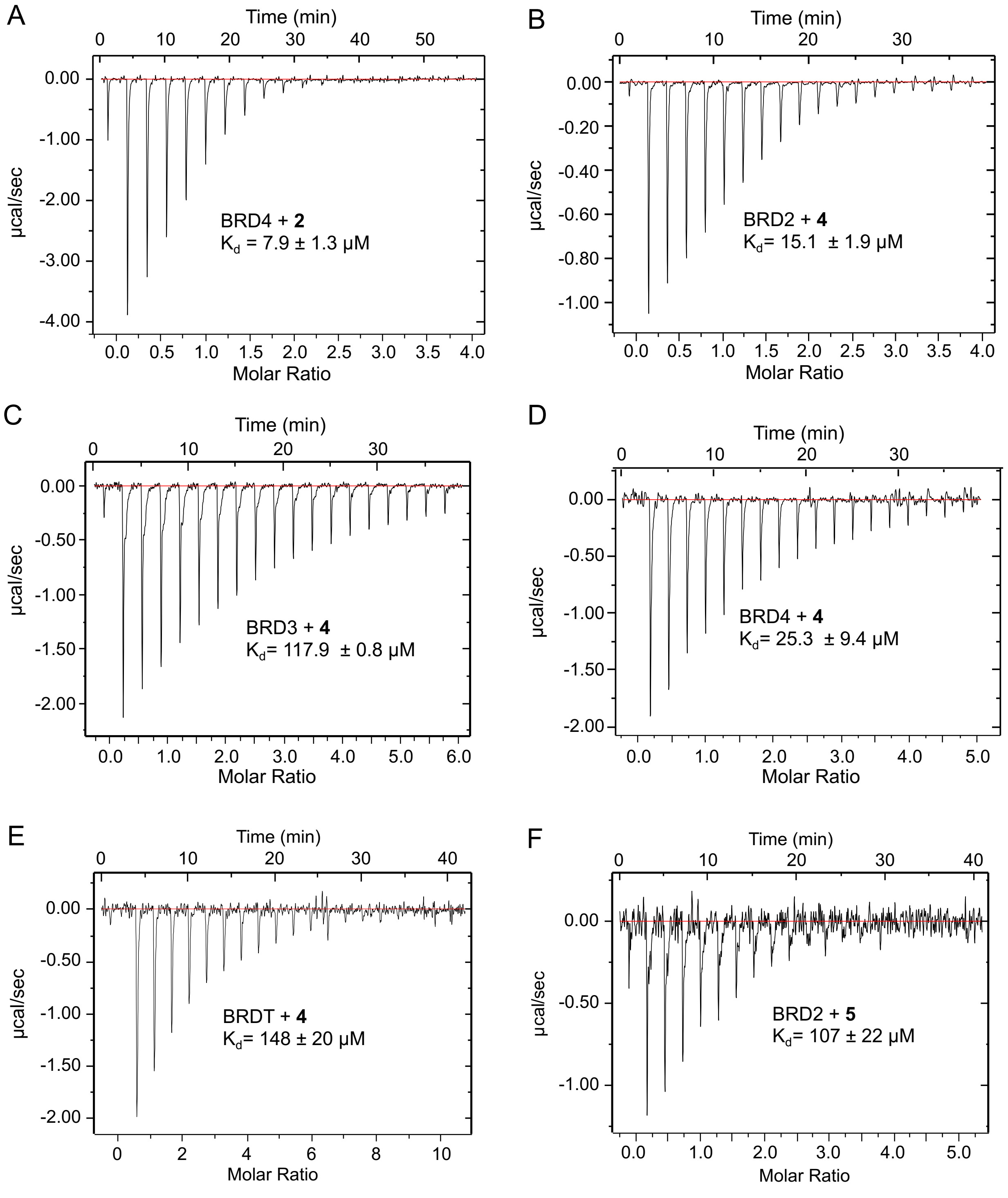


**Supplementary Figure S7**: Binding isotherms of wild type BET BD1 proteins with Peptides **2-13:** (A) WT BRD4-BD1 with H4Kac4 **2**; (B-E) hnRNPK **4** with BRD2, 3, 4, and T, respectively; (F) PD1A1 **5** with BRD2.

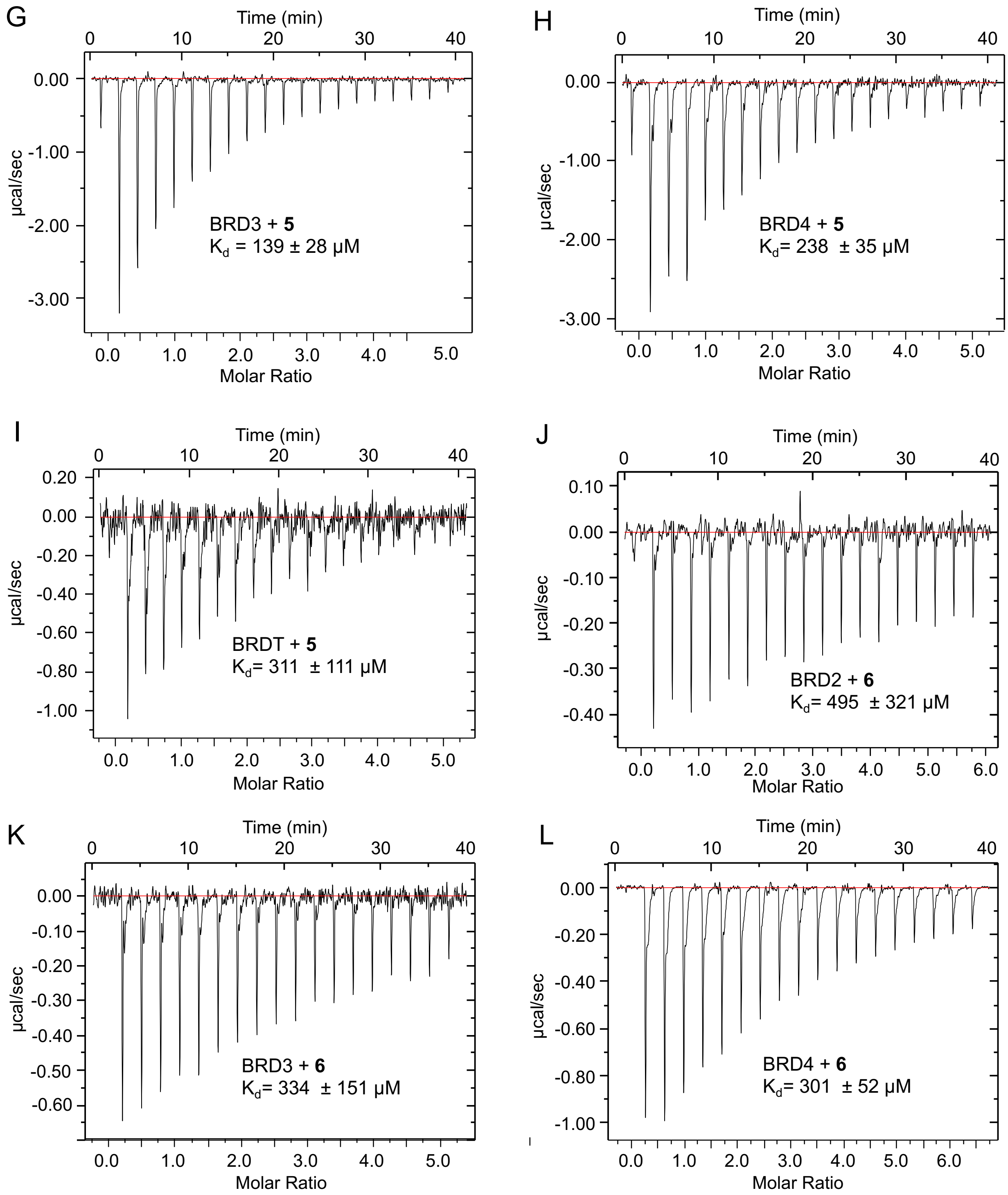


**Supplementary Figure S7 cont'd**: Binding isotherms of wild type BET BD1 proteins with peptides **2-13:** (G-I) PD1A1 **5** with BRD3, 4, and T, respectively; (J-L) PD1A4 **6** with BRD2, 3, and 4, respectively.

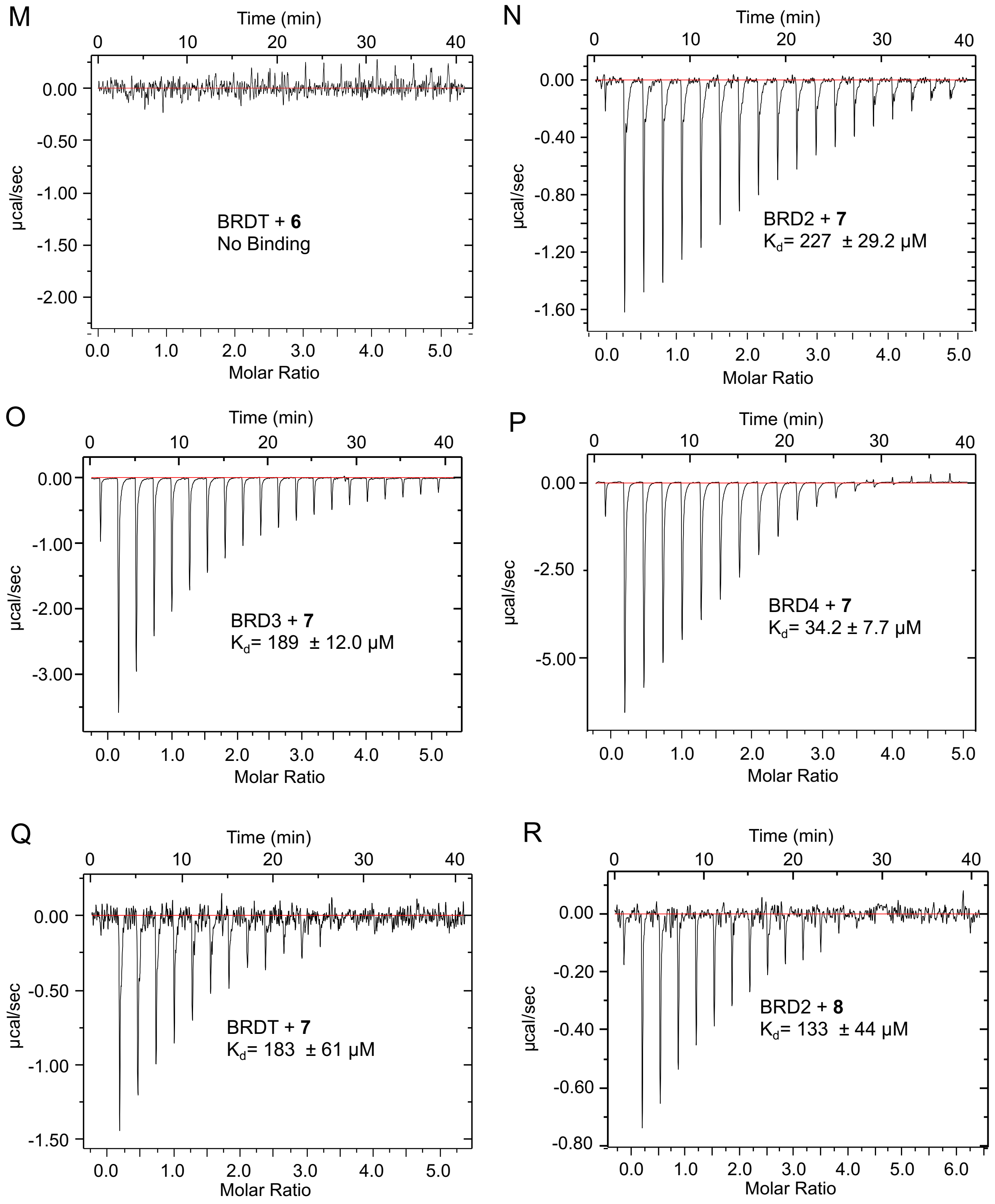


**Supplementary Figure S7 cont'd**: Binding isotherms of wild type BET BD1 proteins with Peptides **2-13:** (M) PD1A4 **6** with BRDT; (N-Q) BCLAF1 **7** peptide with BRD2, 3, 4, and T, respectively; (R) SHMT **8** peptide with BRD2.

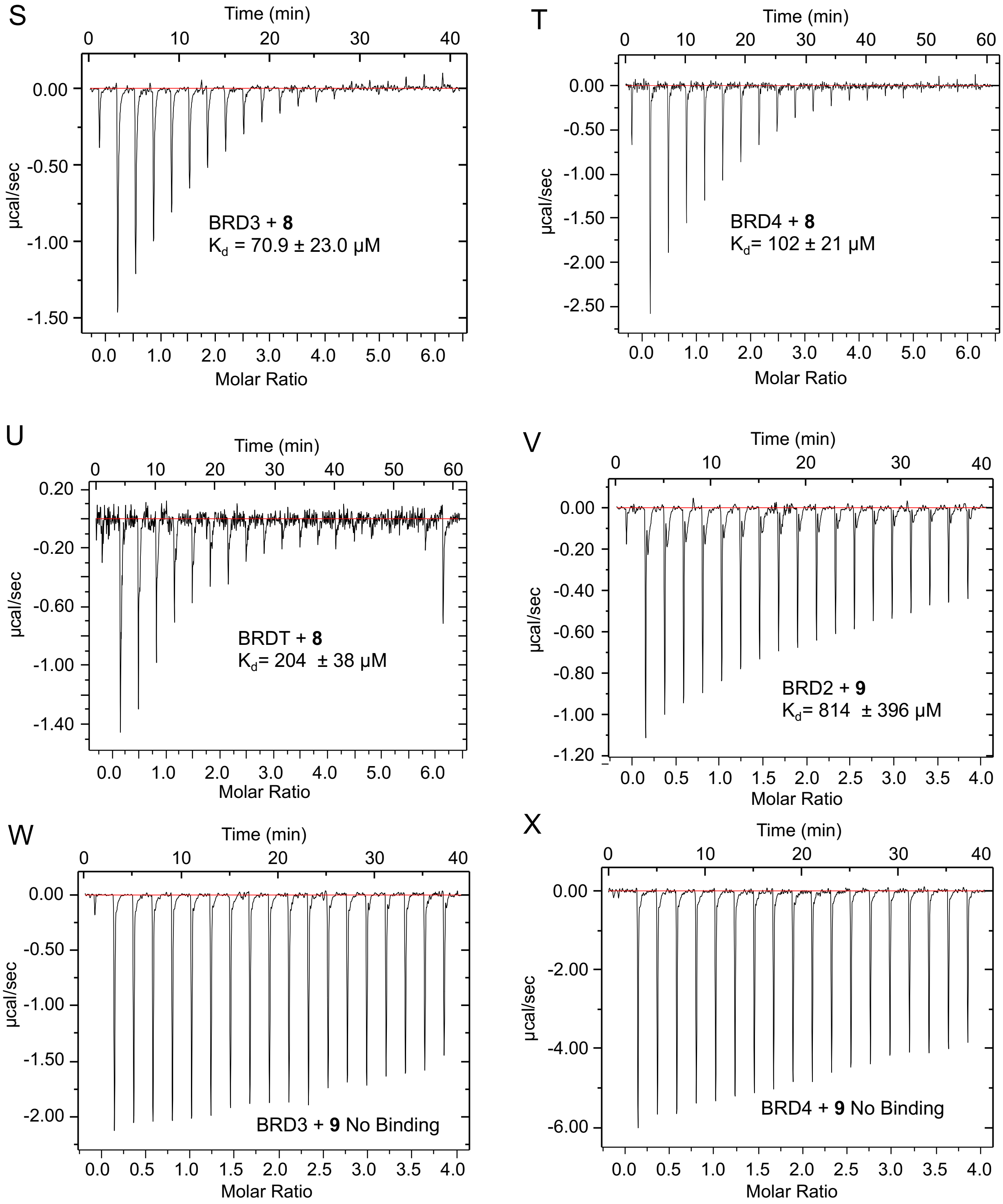


**Supplementary Figure S7 cont'd**: Binding isotherms of wild type BET BD1 proteins with Peptides **2-13:** (S-U) SHMT **8** peptide with BRD3, 4, and T, respectively; (V-X) PCNA **9** peptide with BRD2, 3 and 4.

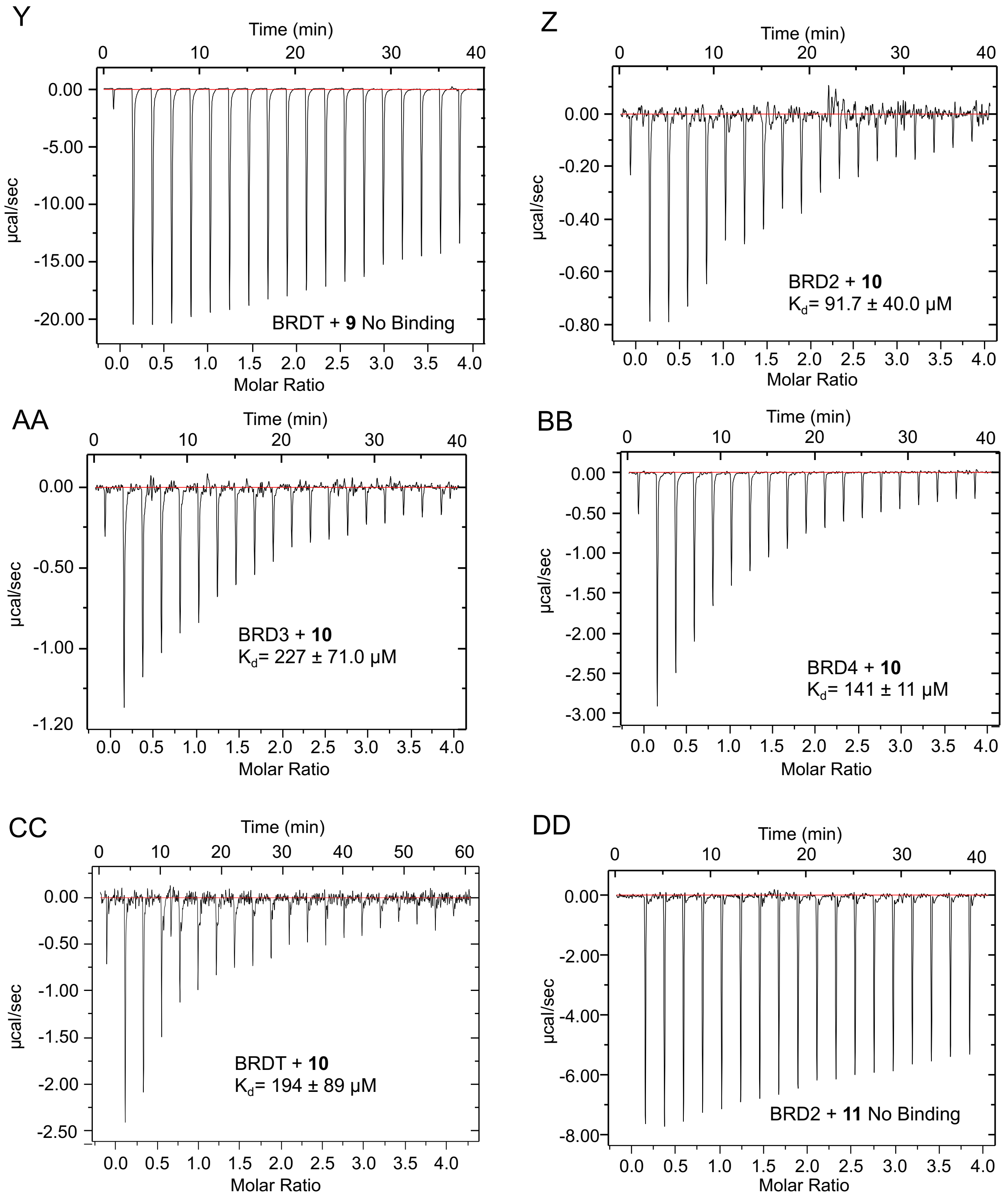


**Supplementary Figure S7 cont'd**: Binding isotherms of wild type BET BD1 proteins with Peptides **2-13:** (Y) PCNA **9** peptide with BRDT; (Z-CC) ILF3 **10** peptide with BRD2, 3, 4, and T, respectively; (DD) 14-3-3Z **11** peptide with BRD2.

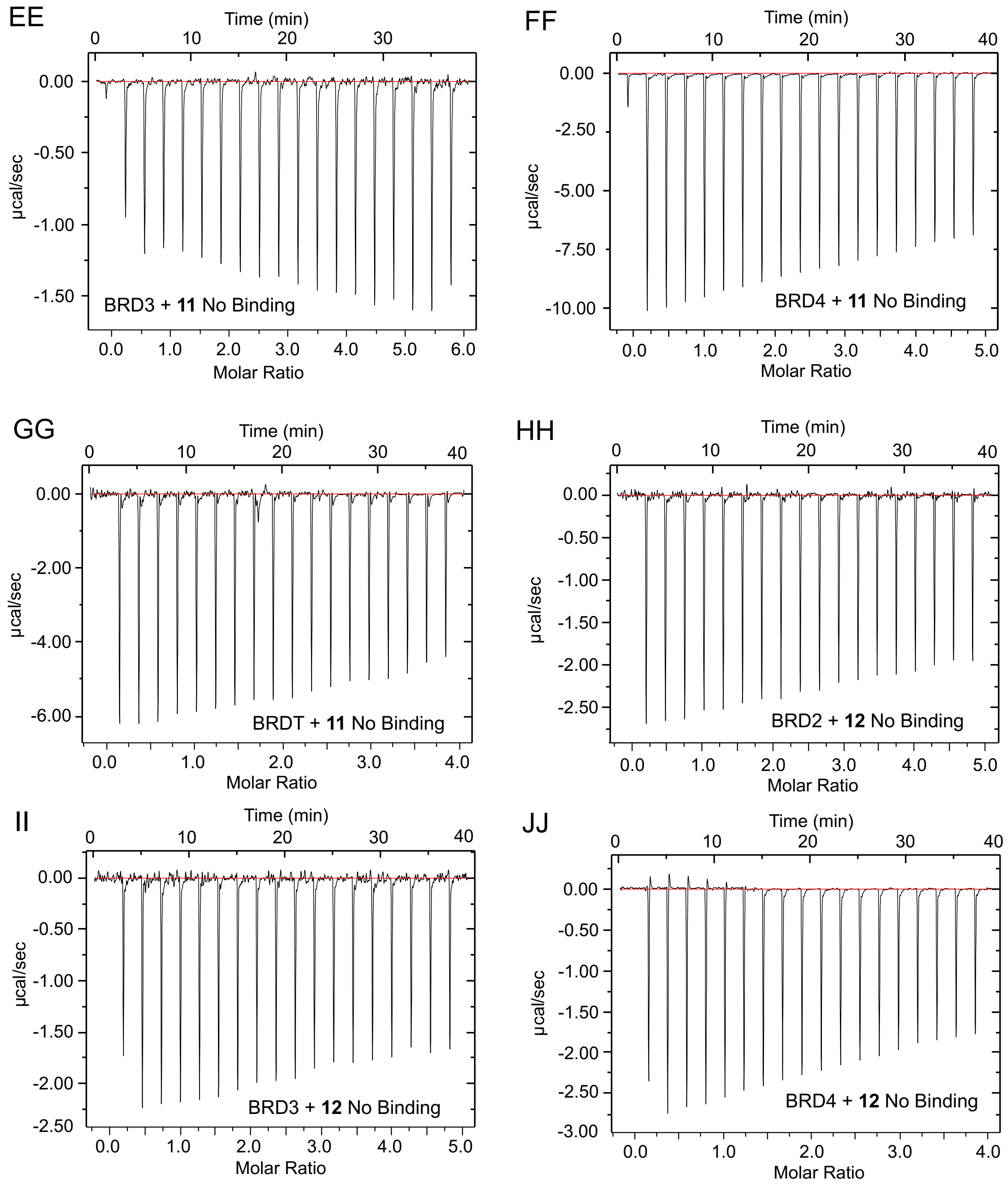


**Supplementary Figure S7 cont'd**: Binding isotherms of wild type BET BD1 proteins with Peptides **2-13:** (EE-GG) 14-3-3Z **11** peptide with BRD2, 3, 4, and T, respectively, (HH-JJ) DHX15 **12** peptide with BRD2, 3 and 4, respectively.

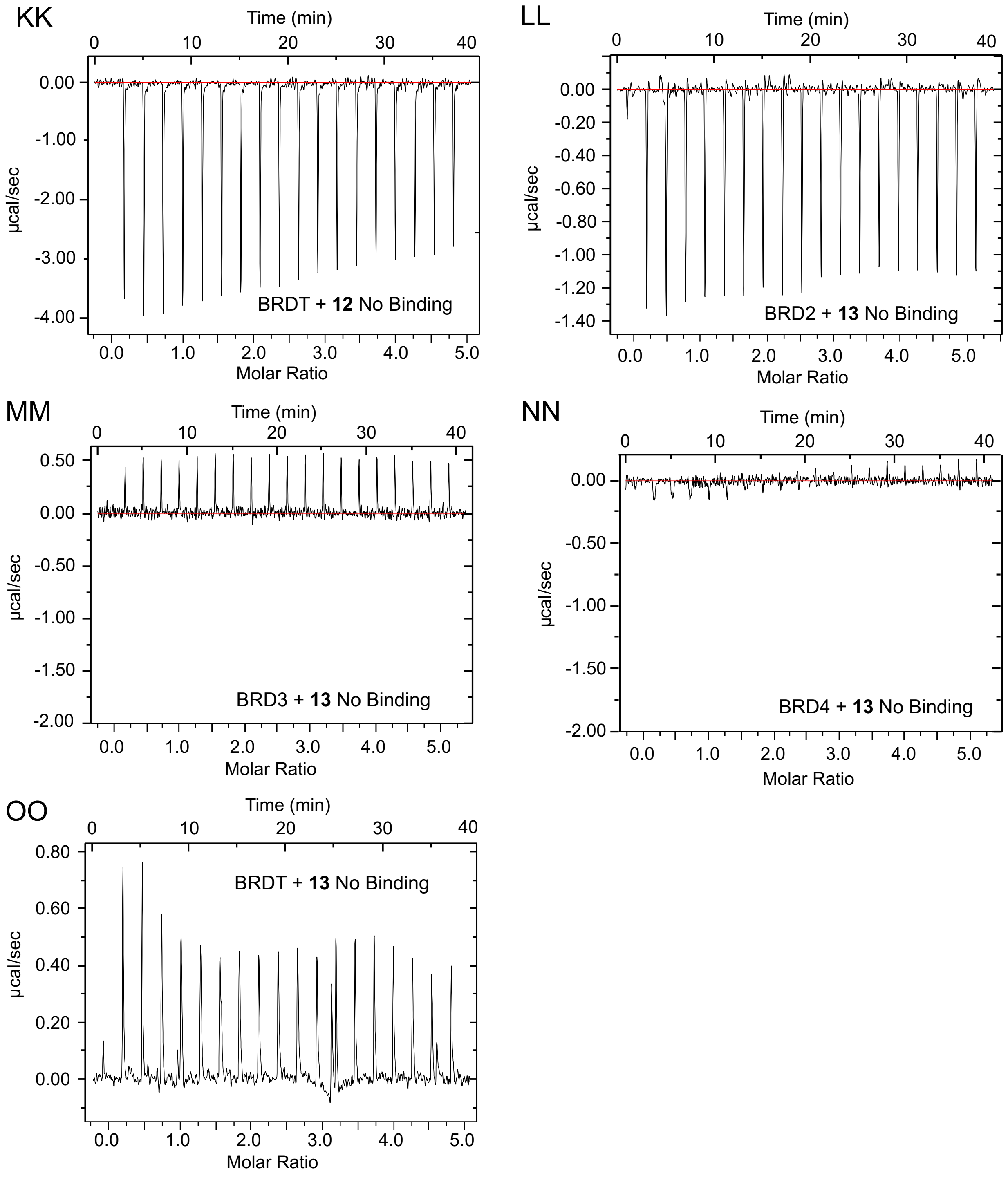


**Supplementary Figure S7 cont'd**: Binding isotherms of wild type BET BD1 proteins with Peptides **2-13:** (KK) DHX15 **12** peptide with BRDT; (LL-OO) XRCC5 **13** peptide with BRD2, 3, 4, and T, respectively.

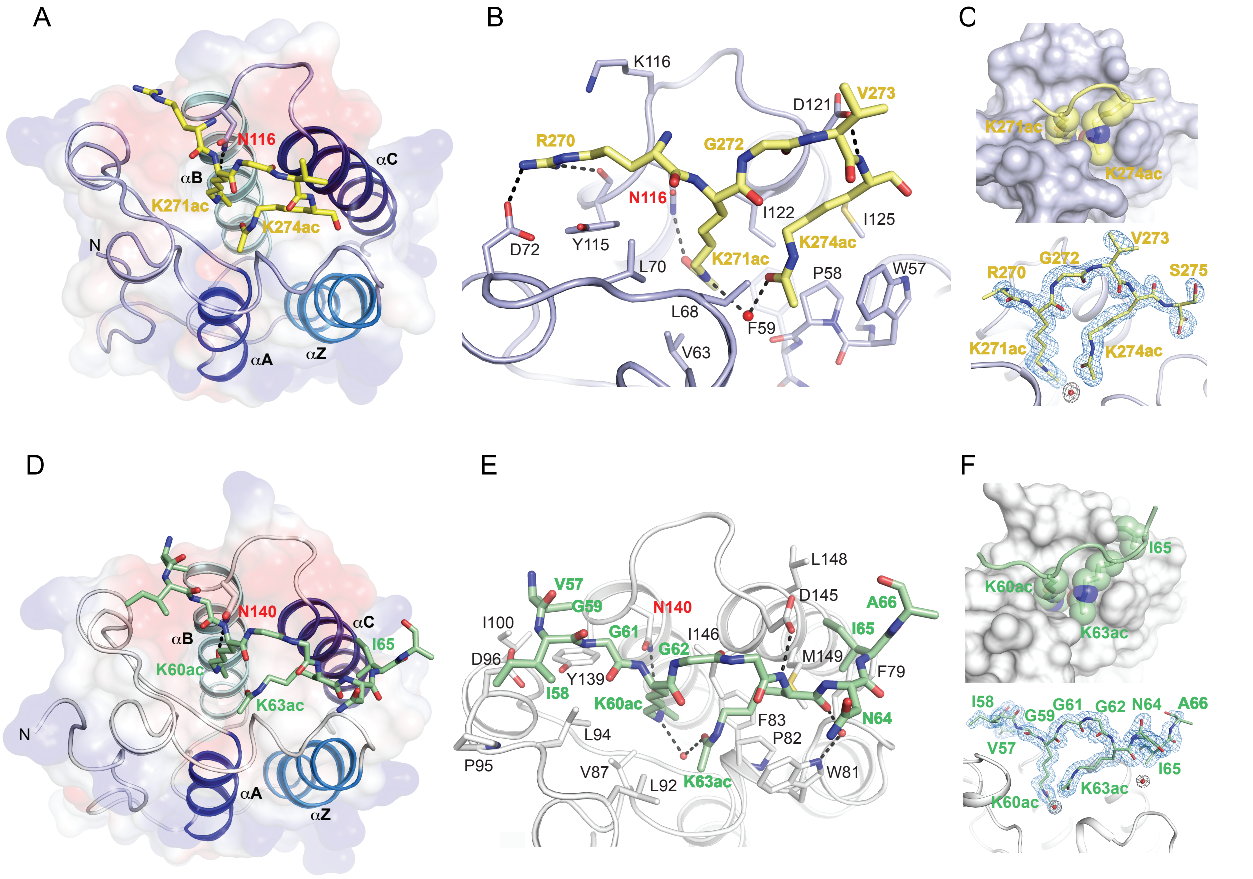


**Supplementary Figure S8**: Structures of BRD3-SHMT and BRD4-hnRNPK complexes. (A-C) A view of BRD3-BD1-SHMT complex containing a continuous 6 amino acid long acetylated (at Lys271 and Lys274) SHMT segment (yellow) (PDB: 7RJL). Interacting side chains in both proteins are depicted in stick representation. Surface representation highlights binding of K271ac and K274ac sidechains of SHMT into the aromatic cage of BRD3. Composite omit maps for the bound acetylated SHMT is shown in blue mesh. Potential hydrogen bonds are denoted with black dashed lines. (D-F) Closeup view of the BRD4-BD1 (light blue) complexed with hnRNPK (green) (PDB: 7RJO). A continuous 9 amino acid long acetylated (at Lys60 and Lys63) hnRNPK peptide that interacts with side chains of BRD4 are shown in stick representation. Surface representation depicts binding of K60ac and K63ac sidechains of hnRNPK into the aromatic cage of BRD4. Electron density from simulated annealing composite omit (5% omission of the model) maps contoured at 1.2 σ (blue mesh) are shown for the bound acetylated protein segment. All figures depicting structure were generated with PyMol.[17]

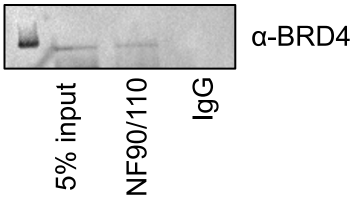


**Supplementary Figure S9**: Immunoprecipitation of BRD4 with NF90/NF110 antibody on Beads followed by Western Blot with BRD4 antibody. IP with control IgG failed to enrich BRD4.

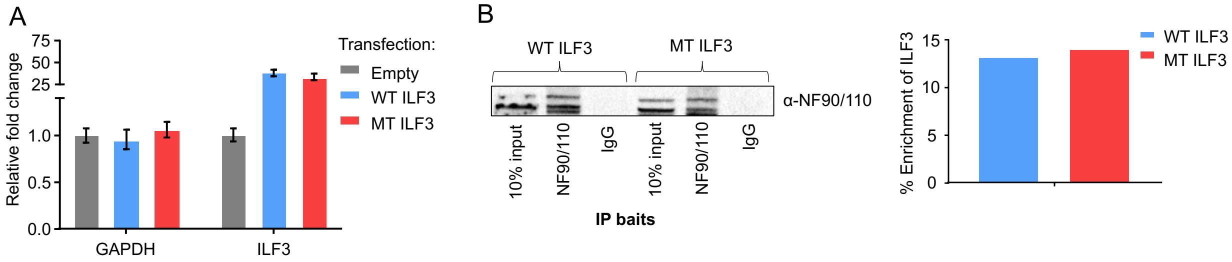


**Supplementary Figure S10**: Assessment of expression and antibody recognition of WT ILF3 and MT ILF3 (K100A/L102A). (A) qRT-PCR with ILF3-specific primers show no statistically significant difference in expression between WT ILF3 and ILF3 K100A/L102A variant. (B) Antibody recognition shows no difference between recognition of WT and recognition of K100A L102A variant at immunoprecipitation (IP) or Western blot (WB) stage. Bar diagram represents quantification of IP based on 10% inputs.

| Protein Name | Amino acid Molecular weight | Noncanonnical Amino Acid Molecular weight AzF or tmdF | Modification | Molecular weight difference from WT | Calculated Molecular weight | Observed (LC-MS) |
|---|---|---|---|---|---|---|
| BRD2-WT | — | — | — | — | 15539.12 | 15537.80 |
| BRD2-L108AzF | Leu-131.17 | 206.20 | L108 to pAzF | 75.03 | 15614.15 | 15613.16 |
| BRD3-WT | — | — | — | — | 17036.52 | 17035.60 |
| BRD3-L68AzF | Leu-131.17 | 206.20 | L68 to pAzF | 75.03 | 17111.55 | 17110.62 |
| BRDT-WT | — | — | — | — | 16614.09 | 16611.25 |
| BRDT-L61AzF | Leu-131.17 | 206.20 | L61 to pAzF | 75.03 | 16689.12 | 16686.16 |

**Supplementary Table S1.** Expected and observed ESI-MS values of BET bromodomains and variants

| Peptide # | Peptide Name | Sequence | Mass |
|---|---|---|---|
| **2** | H4Kac4 | H2N-SGRGK(ac)GGK(ac)GLGK(ac)GGAK(ac)RHRK-CONH2 | 2159 |
| **3** | TAMRA-H4Kac4 | (5/6)-TAMRA-GGGGYSGRGK(ac)GGK(ac)GLGK(ac)GGAK(ac)RHRK-$CONH_2$ | 2963 |
| **4** | hnRNPK | $H_2N$-VIGK(ac)GGK(ac)NIK-$CONH_2$ | 1096 |
| **5** | PD1A1 | $H_2N$-AK(ac)AAGK(ac)LK(ac)A-$CONH_2$ | 983 |
| **6** | PDIA4 | H2N-VTAFK(ac)KGK(ac)LK-CONH2 | 1202 |
| **7** | BCLAF1 | H2N-TAK(ac)TGK(ac)FLKR-CONH2 | 1232 |
| **8** | SHMT | $H_2N$-RK(ac)GVK(ac)S-$CONH_2$ | 756.8 |
| **9** | PCNA | H2N-MSK(ac)ILK(ac)CAG-CONH2 | 1033 |
| **10** | ILF3 | $H_2N$-AK(ac)GLLLK(ac)G-$CONH_2$ | 882 |
| **11** | 14-3-3Z | $H_2N$-MDK(ac)NELVQK(ac)AK(ac)LA -$CONH_2$ | 1613 |
| **12** | DHX15 | $H_2N$-MSK(ac)RHRLD-$CONH_2$ | 1084 |
| **13** | XRCC5 | $H_2N$-LSK(ac)IK(ac)TLF-$CONH_2$ | 1033 |
| **14** | ILF3 no Ac | $H_2N$-AKGLLLKG-$CONH_2$ | 799 |
| **15** | ILF3 L102A | $H_2N$-AK(ac)GALLK(ac)G-$CONH_2$ | 841 |

**Supplementary Table S2:** Sequences and Masses of the peptides used in the current study.

| Protein | Peptide | Protein | Peptide | $K_D$(μM) | N | ΔH | TΔS | ΔG |
|---|---|---|---|---|---|---|---|---|
| | | (μM) | (mM) | | | (kcal/mol) | (kcal/mol) | (kcal/mol) |
| BRD2-BD1 | **2** | 100 | 2 | 27.8 ± 4.0 | 0.931 ± 0.0457 | -9.52 | -3.52 | -6.01 |
| BRD2-BD1 L102AzF | **2** | 100 | 2 | 93.5 ± 19.2 | 0.992 ± 0157 | -14.26 | -8.93 | -5.33 |
| BRD3-BD1 | **2** | 100 | 2 | 15.3 ± 2.1 | 1.02 ± .0292 | -10.78 | -4.44 | -6.34 |
| BRD3-BD1-L68AzF | **2** | 100 | 2 | 14.2 ± 2.1 | 0.800 ± .0291 | -8.45 | -2.06 | -6.39 |
| BRDT-BD1 | **2** | 100 | 2 | 13.7 ± 1.7 | 0.982 ± .0256 | -10.74 | -4.32 | -6.42 |
| BRDT-BD1 L61AzF | **2** | 100 | 2 | 13.5 ± 8.1 | 0.800 ± 0.112 | -4.23 | 2.19 | -6.42 |
| BRD4-BD1 | **2** | 100 | 2 | 7.9 ± 1.3 | 0.952 ± 0.0230 | -12.16 | -5.42 | -6.74 |
| BRD2-BD1 | **4** | 100 | 3 | 15.1 ± 1.9 | 1.29 ± 0.0285 | -2.51 | 3.83 | -6.35 |
| BRD3-BD1 | **4** | 100 | 3.5 | 117.9 ± 8.3 | 1.25 ± 0.0683 | -8.06 | -2.88 | -5.18 |
| BRD4-BD1 | **4** | 100 | 3.5 | 25.3 ± 9.4 | 1.08 ± 0.116 | -5.23 | 0.83 | -6.06 |
| BRDT-BD1 | **4** | 100 | 5 | 148 ± 20 | 1.01 ± 0.191 | -7.37 | -2.32 | -5.05 |
| BRD2-BD1 | **5** | 200 | 5 | 107 ± 22 | 1.19 ± 0.0944 | -2.49 | 2.74 | -5.24 |
| BRD3-BD1 | **5** | 200 | 5 | 139 ± 28 | 0.882 ± 0.131 | -5.17 | -0.08 | -5.09 |
| BRD4-BD1 | **5** | 200 | 5 | 238 ± 35 | 0.896 ± 0.128 | 10.08 | -5.30 | 15.38 |
| BRDT-BD1 | **5** | 200 | 5 | 311 ± 111 | 0.969 ± 0.390 | -4.48 | 0.15 | -4.63 |
| BRD2-BD1 | **6** | 150 | 4 | 294 ± 316 | 4.19 ± 1.13 | -1.07 | 3.60 | -4.67 |
| BRD3-BD1 | **6** | 150 | 4 | 334 ± 151 | 1.46 ± 0.601 | -1.73 | 2.85 | -4.58 |
| BRD4-BD1 | **6** | 150 | 4 | 301.2 ± 5.2 | 1.35 ± 0.236 | -2.58 | 2.07 | -4.64 |
| BRD2-BD1 | **7** | 200 | 5 | 227 ± 29 | 1.37 ± 0.104 | -2.91 | 1.89 | -4.80 |
| BRD3-BD1 | **7** | 200 | 5 | 189 ± 12 | 0.800 ± 0.0556 | -8.84 | -3.92 | -4.92 |
| BRD4-BD1 | **7** | 200 | 5 | 34.2 ± 7.7 | 1.81 ± 0.0646 | -7.59 | -1.70 | -5.89 |
| BRDT-BD1 | **7** | 200 | 5 | 183 ± 61 | 0.800 ± 0.277 | -4.38 | 0.57 | -4.95 |
| BRD2-BD1 | **8** | 200 | 6 | 133 ± 44 | 1.49 ± 0.203 | -0.83 | 4.29 | -5.12 |
| BRD3-BD1 | **8** | 200 | 6 | 70.9 ± 23.0 | 1.32 ± 0.135 | -1.77 | 3.72 | -5.48 |
| BRD4-BD1 | **8** | 200 | 6 | 102 ± 22 | 1.12 ± 0.116 | -3.23 | 2.04 | -5.26 |
| BRDT-BD1 | **8** | 200 | 6 | 204 ± 38 | 0.800 ± 0.176 | -4.76 | 0.11 | -4.87 |
| BRD2-BD1 | **10** | 200 | 4 | 91.7 ± 40 | 1.50 ± 0.189 | -1.22 | 4.09 | -5.31 |
| BRD3-BD1 | **10** | 200 | 4 | 227 ± 71 | 0.800 ± 0.265 | -4.43 | 0.37 | -4.81 |
| BRD4-BD1 | **10** | 200 | 4 | 141 ± 11 | 0.872 ± 0.0481 | -5.73 | -0.65 | -5.08 |
| BRDT-BD1 | **10** | 200 | 4 | 194 ± 89 | 0.800 ± 0.325 | -8.12 | -3.23 | -4.89 |
| BRD4-BD1 | **15** | 200 | 4 | 60.2 ± 10.2 | 0.800 ± 0.0555 | -5.75 | -0.19 | -5.56 |
| BRD4-BD2 | **10** | 150 | 2 | 33.3 ± 6.0 | 0.800 ± 0.0434 | -4.98 | 0.92 | -5.90 |
| BRD4-BD2 | **15** | 150 | 2 | 72.3 ± 8.9 | 0.800 ± 0.0503 | -5.50 | -0.04 | -5.46 |

**Supplementary Table S3:** Thermodynamic parameters measured by isothermal titration calorimetry (ITC) for the binding of the indicated peptides and the BET bromodomains. Conditions are detailed in the experimental section.

**Supplementary Table S4-6:** Analyzed proteomic data for BRD2-L68AzF, BRD3-L108AzF and BRDT-L61AzF. These tables are provided separately.

**Supplementary Table S7:** Crystallization conditions, crystallographic statistics and RCSB accession codes. This table is provided separately.

| GENE | VECTOR | AFFINITY TAG | RESISTANCE | SOURCE |
|---|---|---|---|---|
| BRD2-BD1 | pET | N-6xHis | Ampicillin | VectorBuilder |
| BRD3-BD1 | pNIC28-Bsa4 | N-6xHis | Kanamycin | #65377 Addgene |
| BRD4-BD1 | pNIC28-Bsa4 | N-6xHis | Kanamycin | #38942 Addgene |
| BRD4-BD2 | pNIC28-Bsa4 | N-6xHis | Kanamycin | #38943 Addgene |
| BRDT-BD1 | pNIC28-Bsa4 | N-6xHis | Kanamycin | #38898 Addgene |
| *M. j.* TyrRS-tRNA$^{CUA}$ | pEVOL | None | Chloramphenicol | #31186 Addgene |
| HISTONE H4 | pDEST | None | Ampicillin | Sloan Kettering Institute |
| BRD4 | pcDNA4c | His | Ampicillin | #14441 Addgene |
| BRD4 | pcDNA5/frt to | None | Ampicillin | #90331 Addgene |
| BRD4 N140A | pcDNA5/frt to | None | Ampicillin | VectorBuilder |
| BRD4 N433A | pcDNA5/frt to | None | Ampicillin | VectorBuilder |
| NF90 | pcDNA | None | Ampicillin | ETH Zurich |
| NF110 | pcDNA3.1 | None | Ampicillin | VectorBuilder |
| GCN5 | pEBB | None | Ampicillin | #74784 Addgene |
| CBP | pcDNA3 | None | Ampicillin | #32908 Addgene |

**Supplementary Table S8.** List of the genes used in the current study. The expression vector, affinity tag for protein purification, antibiotic resistance, and source are provided.

| MUTATION | FORWARD PRIMERS |
|---|---|
| BRD2-BD1-L108TAG | 5'-CCTGTGGATGCTGTCAAATAGGGTCTACCGGATTATCAC-3' |
| BRD3-BD1-L68TAG | 5'-CCGTGGACGCAATCAAATAGAACCTGCCGGATTATC-3' |
| BRD4-BD1-L92TAG | 5'-GTGGATGCCGTCAAGTAGAACCTCCCTGAT-3' |
| BRDT-BD1-L61TAG | 5'-GTGGATGCTGTGAAATAGCAGTTGCCTGAT-3' |
| H4-K5C | 5'-GTCTGGTCGTGGTTGCGGTGGTAAAGGT-3' |
| ILF3_K100A_L102A_Fwd | 5'-GTGGGCCTGGTGGCAGCGGGCGCGCTACTCAAGGGG-3' |

**Supplementary Table S9.** List of primers designed for site-directed mutagenesis. Reverse primers used are the reverse-complement to the given forward primers.

| MUTATION | FORWARD PRIMERS |
| --- | --- |
| Survivin_ Fwd | CTGCCTGGCAGCCCTTT |
| Survivin_Rev | CCTCCAAGAAGGGCCAGTTC |
| B-Actin_ Fwd | CTCCTTAATGTCACGCACGAT |
| B-Actin_ Rev | CATGTACGTTGCTATCCAGGC |
| GAPDH_ Fwd | GTGGTCTCCTCTGACTTCAACAGC |
| GAPDH_ Rev | GAGCTTGACAAAGTGGTCGTTGAG |
| BRD4_Fwd | ACCTCCAACCCTAACAAGCC |
| BRD4_Rev | TTTCCATAGTGTCTTGAGCACC |
| ILF3_Fwd | AGCATTCTTCCGTTTATCCAACA |
| ILF3 _Rev | GCTCGTCTATCCAGTCGGAC |
| Survivin_TATA_Fwd | TTAACCGCCAGATTTGAATCGCGG |
| Survivin_TATA_Rev | CCAAGAAGGGCCAGTTCTTGAATG |

**Supplementary Table S10.** List of primers used for qRT-PCR and ChIP.[18-19]
